# Macro-environment strongly interacts with warming in a global analysis of decomposition

**DOI:** 10.1101/2024.04.03.587921

**Authors:** Sarah Schwieger, Ellen Dorrepaal, Matteo Petit Bon, Vigdis Vandvik, Elizabeth le Roux, Maria Strack, Yan Yang, Susanna Venn, Johan van den Hoogen, Fernando Valiño, Haydn J.D. Thomas, Mariska te Beest, Satoshi Suzuki, Alessandro Petraglia, Isla H. Myers-Smith, Tariq Muhammad Munir, Anders Michelsen, Jørn Olav Løkken, Qi Li, Takayoshi Koike, Kari Klanderud, Ellen Haakonsen Karr, Ingibjörg Svala Jónsdóttir, Robert D. Hollister, Annika Hofgaard, Ibrahim A. Hassan, Wang Genxu, Nina Filippova, Thomas W. Crowther, Karin Clark, Casper T. Christiansen, Angelica Casanova-Katny, Michele Carbognani, Stef Bokhorst, Katrín Björnsdóttir, Johan Asplund, Inge Althuizen, Rocío Alonso, Juha Alatalo, Evgenios Agathokleous, Rien Aerts, Judith M. Sarneel

## Abstract

Empirical studies worldwide show substantial variability in plant litter decomposition responses to warming, leaving the overall impact of climate change on this process uncertain. We conducted a meta-analysis of 109 experimental warming studies across seven continents, utilizing natural and standardized plant material, to assess the overarching effect of warming on decomposition and identify potential moderating factors.

Warming influences decomposition differently across macro-environmental gradients of moisture and temperature. Negative warming effects on decomposition in warmer, low-moisture areas were counterbalanced by the positive, though not significant, warming effects in colder areas, resulting in an overall non-significant effect. We determine that at least 5.2 degrees of warming is required for a significant increase in decomposition. This is particularly relevant given the past decade’s global warmth in higher latitudes, holding a significant proportion of terrestrial carbon. Low-quality plant litter was more sensitive to warming. Therefore, future vegetation changes toward low-quality, temperature-sensitive plants could increase carbon release and reduce the net supply of stored organic matter in the soil by increasing the decomposition of low-quality litter with warming. Our findings emphasize the connection between warming responses, macro-environment, and litter characteristics, refining predictions of climate change’s consequences on key ecosystem processes and its contextual dependencies.

## Introduction

The temperature sensitivity of plant litter decomposition is key to understanding future nutrient and carbon cycling, as any change in decomposition resulting from climate warming may alter nutrient availability, plant growth, and carbon storage in terrestrial ecosystems (Swift *et al*. 1979; Bardgett *et al*. 2013). Carbon modelling (e.g., Jenkinson *et al*. 1991; Schimel *et al*. 1994; Davidson *et al*. 2000; Kirschbaum 2004; Knorr *et al*. 2005), kinetic theory (e.g., Davidson and Janssens 2006), and laboratory incubations (e.g., Kirschbaum 1995; Rey *et al*. 2008; Conant, Steinweg, *et al*. 2008) show that decomposition rates increase with increasing temperature. However, site-specific empirical field studies reveal significant variations in the responses of decomposition rates to experimental warming, ranging from increased (Shaw and Harte 2001; Luo *et al*. 2010; Berbeco *et al*. 2012; Henry and Moise 2015; Ren *et al*. 2018) or no effect (Zaller *et al*. 2009; Cheng *et al*. 2010; Moise and Henry 2014; Gong *et al*. 2015), to decreased decomposition under warming (Christiansen *et al*. 2017; Romero-Olivares *et al*. 2017; Hong *et al*. 2021). Results from these site-specific studies pose a challenge when attempting to generalize how ongoing climate change will affect nutrient and carbon cycling across ecosystems. Here, we synthesise the latest available results from *in situ* experimental warming studies across terrestrial ecosystems worldwide that measured the mass loss of plant litter. We combine these results with the implementation of a globally distributed, standardised decomposition assay to improve our understanding of how and where climate warming may affect plant litter decomposition, and to identify potential moderating factors.

Litter decomposition is a complex process, involving the biological (i.e., microbial, enzymatic mineralisation), chemical and physical transformation and breakdown of organic matter (Swift *et al*. 1979; Bardgett *et al*. 2008; Kirchman 2018), including leaching of soluble chemicals into the soil (Coûteaux *et al*. 1995). Warming can directly stimulate microbial and enzymatic activity (Bardgett *et al*. 2008), as well as leaching (Lind *et al*. 2022), and thus increase decomposition rates. However, it is widely recognised that warming is likely to increase *in situ* decomposition rates more where there is sufficient soil moisture (Aerts 2006). Biomes inherently vary in temperature and precipitation, irrespective of climate change, contributing to the diverse decomposition responses observed in *in situ* experiments globally. Consequently, projected increases in temperature and changes in precipitation changes may significantly alter decomposition rates (Zhang *et al*. 2008; Conant *et al*. 2011; Wu *et al*. 2011; Djukic *et al*. 2018; Joly *et al*. 2023).

While different types and degrees of climate change may thus have different effects on plant litter decomposition, the type and intensity of climate change is not uniform across the globe (IPCC 2021). Such spatial differences in climate change, combined with differences in local environmental conditions, and potential interactions between local environmental conditions and climate change, are likely to lead to spatial variation in decomposition responses to warming. For example, cold temperatures tend to inhibit decomposition, creating huge carbon stores in high latitude soils (Schuur *et al*. 2008; Tarnocai *et al*. 2009). At the same time, decomposition at low temperatures is particularly sensitive to small changes in temperature (Kirschbaum 1995; Chen *et al*. 2015). Therefore, climate warming is expected to increase decomposition more strongly in colder high-latitude and high-altitude regions, creating a positive carbon-climate feedback loop where warming releases greenhouse gasses from these carbon rich soils (Jenkinson *et al*. 1991; Kirschbaum 1995, 2000; Cox *et al*. 2000; Fenner and Freeman 2011). In other places, climate warming is predicted to increase the frequency and intensity of droughts, which could in turn reduce decomposition by limiting the biological activity of decomposer organisms, as has been shown in temperate grasslands (Vogel *et al*. 2013; Walter *et al*. 2013). Therefore, in warmer systems and systems with high variability in precipitation (e.g., savannahs), the warming response is likely to depend strongly on concurrent moisture conditions (Aerts 1997; Seres *et al*. 2022). Our study should lead to a better understanding of the interaction between the prevailing macro-environmental conditions and the changes in decomposition caused by warming. This will help us to better predict the consequences for carbon and nutrient cycling in terrestrial ecosystems. Since the initial conditions and the intensity of climate change vary greatly around the world, the responses of plant decomposition to warming will also be very different. Therefore, our study aims to identify recognisable patterns in decomposition responses to warming under different macro-environmental conditions.

There is increasing evidence that litter quality (i.e., the chemical characteristics of the decomposing material) may control the temperature sensitivity of litter decomposition (Bosatta and Ågren 1999; Fierer *et al*. 2005; Davidson and Janssens 2006; Conant, Drijber, *et al*. 2008; Suseela *et al*. 2013). Despite the complex and diverse chemical make-up of plant litter, comparisons of its relationship to decomposition across plant species on a global scale often rely on assessing chemical composition through factors such as carbon to nitrogen (C:N) ratios (Aerts 1997; Prescott 2010), plant functional types (e.g., trees, shrubs, mosses, graminoids) (Chapin *et al*. 1996; Dorrepaal *et al*. 2005), or plant organs (e.g., shoots, leaves, roots) (Freschet *et al*. 2012; Xia *et al*. 2015). Additionally, decomposability, a measure of the decomposition rate under ambient conditions, provides information about the pace at which plant litter breaks down but can also serve as a proxy for assessing litter quality (Cornelissen *et al*. 2004; Freschet *et al*. 2012). Litter with a low C:N ratio (e.g., of forbs, leaves) generally decomposes faster than litter with a high C:N ratio (e.g., of mosses, trees and shrubs, roots) due to its higher quality compared to materials rich in lignin, tannin and other complex carbohydrates (Swift *et al*. 1979; Fontaine *et al*. 2004; Kirchman 2018). Litter with a high C:N ratio or a low decomposability is thought to be more temperature sensitive, and warming could therefore disproportionately accelerate the decomposition of such litter compared with a low C:N ratio litter or a high decomposability (Biasi *et al*. 2005; Davidson and Janssens 2006; Conant, Steinweg, *et al*. 2008). Clarifying the role of litter quality in mediating warming effects on decomposition, and its how it displays across macro-environmental gradients, is pivotal for addressing the variability observed in *in situ* studies. This understanding is crucial for predicting changes in nutrient and carbon cycling with warming across diverse plant communities. Notably, as warming alters plant community composition in certain regions, for example, arctic tundra and alpine areas, it becomes essential to comprehend how this shift influences litter composition and quality (Elmendorf *et al*. 2012; Pearson *et al*. 2013; Munir *et al*. 2017).

In this study, we aim to quantify the effect of experimental warming on plant litter decomposition across a global range of ecosystems and environmental regions, and to identify the context-dependencies of variable warming effects at a global scale. To this end, we assessed the influence on the warming effect on decomposition of (1) macro-environmental regions (i.e., site-level environmental conditions), (2) experimentally induced changes in micro-environment (i.e., plot-level temperature and moisture changes with warming), and (3) litter quality within macro-environmental regions. We expected that:

i. the macro-environmental region is a key determinant of the effect of warming on decomposition. In temperature-limited systems, we expect a higher sensitivity and an increase in decomposition with warming, whereas in moisture-limited systems, we expect a lower sensitivity to warming and a decrease in decomposition.
ii. A stronger warming will proportionally increase decomposition if warming does not limit moisture.
iii. Litter quality modulates the effect of warming on decomposition with lower quality litter being more sensitive to warming than high quality litter.

To test these hypotheses, we conducted a global meta-analysis, examining 109 datasets with experimental setups comprising 637 paired (i.e., warmed and ambient) observations of decomposition of plant litter under ambient conditions vs. experimental warming. These datasets were obtained from *in situ* warming experiments that either decomposed natural local plant species litter (52 paired studies, sourced from published literature) or two standardised plant litter, green and rooibos tea (57 paired experiments each, from unpublished, primary research data). This comprehensive analysis allowed us to quantify the effects of warming on natural litter decomposition, where plant community specific local litter is decomposed at the site of origin, while the use of standardised plant litter allowed filter out home effects and just be able to assess and compare the role of environmental factors across locations.

## Methods

In this meta-analysis, we combined two global datasets. First, we extracted data from the 52 published studies that measured decomposition responses of natural litter to experimentally imposed higher temperatures. Further, we buried rooibos and green tea as standardised plant litter in 57 warming experiments (Keuskamp *et al*. 2013). Whereas the natural litter data mainly covers the United States, Western Europe and China, the standardised plant litter decomposition data ranges from higher latitudes as well as the Mediterranean and a few sites in the southern hemisphere (Figure 1).

**Figure 1.**
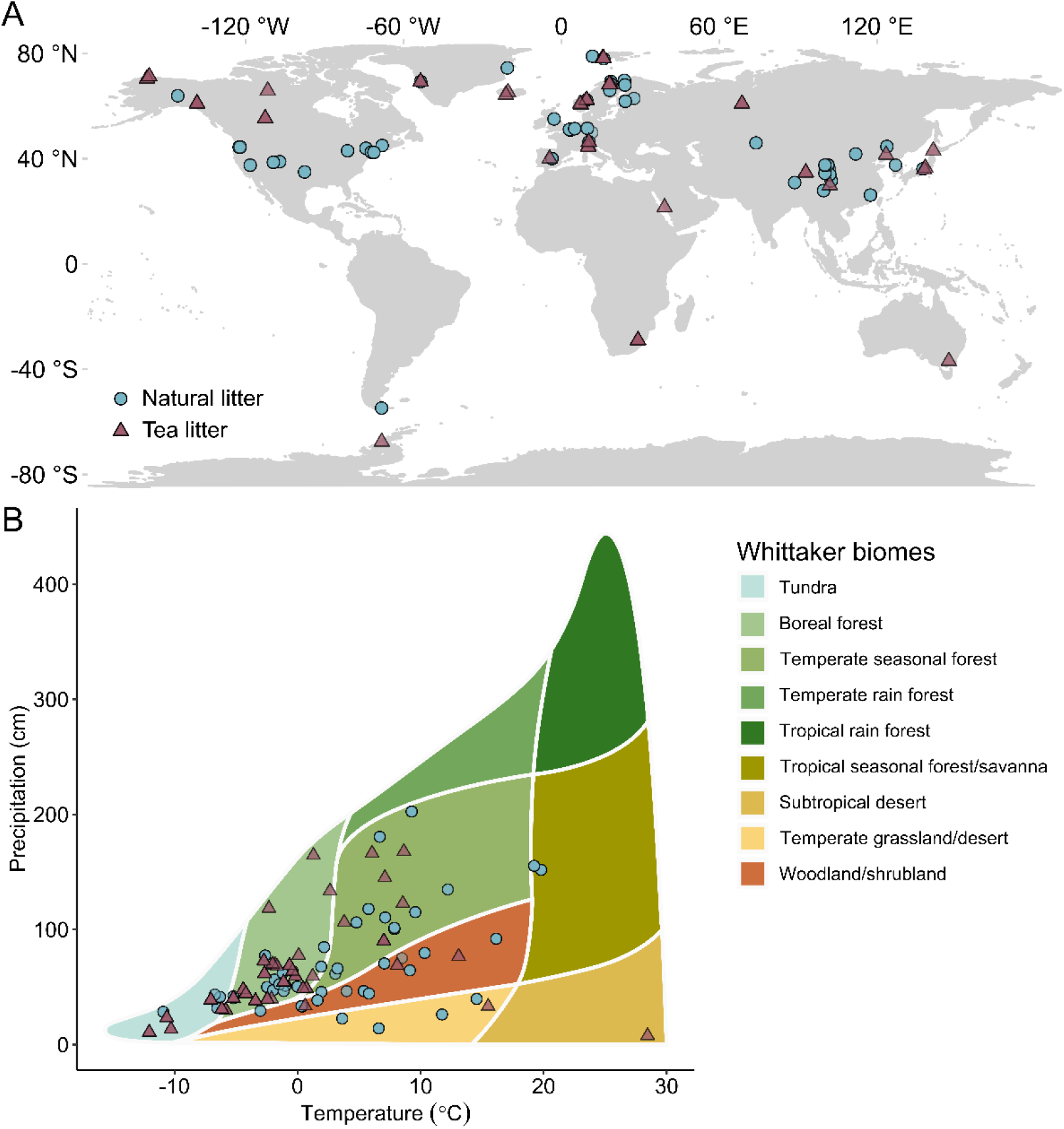
(**A**) Map and (**B**) Whittaker biome diagram showing the location of the 52 published studies of natural litter decomposition in warming experiments (blue circles) and the location of the 57 open-top chamber experiments where we deployed tea as a standardised plant litter to assess decomposition response to warming (purple triangles) used in this meta-analysis. Our study does not cover temperate and tropical rainforests.

### Data collection

#### Literature data on natural plant litter decomposition

We conducted an extensive literature survey for peer-reviewed publications in the ISI Web of Science database (http://apps.webofknowledge.com/) on September 1^st^ 2023. We used *(warming OR heat* OR OTC OR open top chamber*) AND (litter* OR litter bag) AND (decomposition OR mass loss)* as search criteria, which returned 1184 studies (Figure S1). We considered terrestrial field studies that compared litter decomposition (mass loss and decomposition rate) under experimentally increased temperatures (methods found in our search were open-top chambers, heating cables, infrared heaters, sunlit controlled-environment chambers, UVB filter films, open-topped polythene tents, and closed-top chambers) and ambient conditions. From the 60 studies that met our criteria, we extracted mean values, sample sizes and measures of variation (i.e., standard errors or standard deviations) for decomposition (i.e., decomposition rates, absolute and relative mass loss, remaining mass of plant material). We contacted the corresponding authors to obtain access to the raw data for studies that did not report them and had to exclude 8 studies due to insufficient reporting. This resulted in 52 studies used for the meta-analysis (Table S1). Whenever warming was applied in factorial combination with one or more additional treatments (e.g., warming and plant species removal), we extracted the parameters of interest for the warming treatment only, together with the ambient control. If the litter was incubated at different time steps, each time step was used as an independent data point. We thus extracted a total of 523 paired (ambient vs warmed) data points from the 52 studies, either directly from the text or tables or from figures using the software WebPlotDigitizer (v. 4.6, Rohatgi 2021). When decomposition was reported as remaining mass of plant material, the latter was transformed into mass loss.

We extracted coordinates of each study location (Figure 1A), the incubation duration of the litter (from 14 days to 4.9 years, standardised to days), the mesh size of the litter bags (from 0.02 to 5 mm), the position of incubation (i.e., if litter bags were put on the soil surface or buried below ground), the plant species and thus the plant functional type (i.e., forb, nonvascular, graminoid, woody species), and the plant organ type (i.e., leaf, shoot or root). For 32 studies, we also extracted litter C:N ratio reported by the researchers (ranging from 12 to 201). All reported values were from single species, with the only exception being two studies on root decomposition, which included a mixture of grass species. Yet, as all the species in these samples were indeed within the graminoid functional type, we included these studies in the meta-analysis. For each study, we extracted the duration of the warming experiment prior to incubation start (from first year to 23 years). The warming method was classified as Heating cables (number of studies n=11), Infrared heaters (n=17), and Open-top chambers (n=19), with ‘Other methods’ including Sunlit controlled-environment chambers (n=1), UVB filter films (n=1), Open-topped polythene tents (n=2) and Closed-top chambers (n=1).

#### Standardised plant litter data from open-top chamber warming experiments

Following the standard Tea Bag Index protocol (Keuskamp et al., 2013), green (*Camellia sinensis*; EAN no.: 8 722700 055525) and rooibos (*Aspalathus linearis*; EAN no.: 8 722700 188438, Lipton, Unilever) tea bags with woven nylon mesh (0.257 mm), were buried at a depth of 8 cm and at a distance of at least 15 cm from each other in open-top chambers (OTC) and controls at 57 locations (Figure 1, Table S2). The incubations covered one growing season (82 ± 18 days; mean ± SD), that is, from May/June 2016 to August/September 2016 in the northern hemisphere and from January 2017 to March 2017 in the southern hemisphere. For two sites in Japan (i.e., JPN_1 and JPN_3, Table S2), tea bags were incubated from July to October 2012. Retrieved bags were cleaned of adhering soil and roots. The mass of the remaining tea was determined after drying it in an oven at 60-70 °C for at least 48 h. To align with the literature data, we calculated treatment means of mass loss, sample sizes and standard deviations for each experiment/GPS location.

#### Explanatory macro-environmental drivers

We obtained map-based environmental data based on the geographical locations of the study sites to identify macro-environmental factors that may influence the response of decomposition to warming. We used 48 environmental layers reflecting major gradients in climate, soil, vegetation, and topographic variables as covariates in our analysis (Table S3).

Due to the occurrence of many confounding variables, we summarised the macro-environmental variation across studies with a Principal Component Analysis (PCA; Table S3). The first principal component (PC 1) was strongly positively correlated with temperature-associated variables and negatively correlated with soil organic carbon (SOC) and explained 26.9 % of the total variance (Figure 3A, Table S3). The second component (PC 2) correlated positively with precipitation-associated variables and explained 18.1% of the total variance (Figure 3A, Table S3). The third PC axis was not considered as it described negligible amounts of the variation (4.2%). In our dataset, the range of annual mean temperature was −12 to 28 °C, annual precipitation was 78 to 2100 mm, and soil saturated water content was 42 to 81 %. We created four ‘macro-environmental classes’ based on the origin of the PC1 and PC2 variables as a separation line. These four ‘macro-environmental classes’ were described as following: (1) high temperatures and high precipitation (number of effect sizes *k*=156), (2) high temperatures and low precipitation (*k*=170), (3) low temperatures and high precipitation (*k*=156), and (4) low temperatures and low precipitation (*k*=155) (Figure S2, Table S4).

#### Explanatory micro-environmental drivers altered by experimental warming

For both datasets (i.e., natural and standardised plant litter), we collected available data on the actual degree of warming, i.e., the mean absolute temperature difference between the warmed and ambient control, as well as soil moisture in warming and control treatments when available. The degree of warming included air or soil temperature measures, depending on whether the litter was incubated on the soil surface or below ground, respectively. We calculated relative change in soil moisture with warming according to:

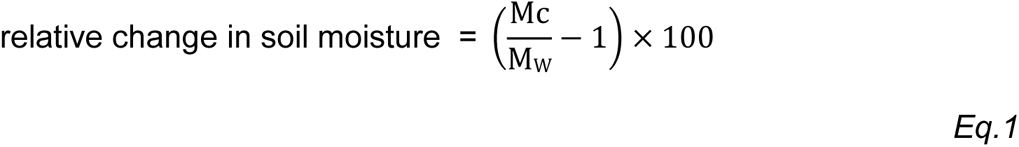

where M_C_ and M_W_ are soil moisture in control and warming treatment, respectively. Positive and negative values indicate drier and wetter conditions under warming than under ambient conditions, respectively.

#### Litter quality

We focused on three different, frequently used characterisations of litter qualities: the C:N ratio before decomposition, reported in the initial studies, the decomposability measured as decomposition rate under ambient condition (i.e., standardised to mass loss in % d^-1^) (Cornelissen *et al*. 2004; Freschet *et al*. 2012), and plant functional type (Dorrepaal *et al*. 2005). We categorised the plant species into four different plant functional types (sensu Chapin *et al*. 1996), forbs (number of studies n=7), graminoids (i.e., grasses and sedges, n=28), woody species (i.e., shrubs and needle-leaved and broad-leaved trees, n=27), and nonvascular (i.e. mosses, n=4; lichens, n=1). For graminoids and woody species, we were able to further specify litter type into aboveground (i.e., shoots and leaves of graminoids, n=25; broadleaves and needles of woody species, n=25) and below-ground plant organs (i.e., roots of graminoids, n=6; and root of woody species, n=2).

### Data analysis

In order to evaluate the relative effect of experimental warming on decomposition, we used Hedges’ g, which is a standardised mean difference (SMD) calculated by dividing the difference between the mean mass loss in the warming treatment (*x̅*_1_) and control (*x̅*_2_) by the pooled standard deviation (Hedges 1981).

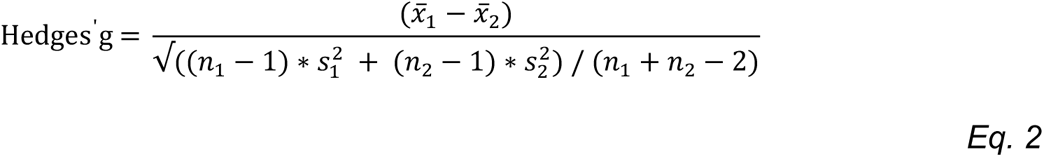

where n_1_ and n_2_ are sample size, and s_1_^2^ and s_2_^2^ are the sample variance of the warming treatment and the control, respectively. Therefore, a SMD larger than one indicates that warming enhanced decomposition, while a SMD lower than one indicate that warming decreased decomposition. By using the SMD as a measure of effect size, we were able to synthesize data measured on different scales or units (e.g. mass loss vs. decomposition rate), while still accounting for the precision (variance) of the measurement. We used the *escalc()* function in the R package METAFOR (v.4.0-0; Viechtbauer 2010) to derive SMDs and corresponding 95% confidence intervals (CI) from all paired (i.e., warmed/control) mean values, sample sizes and standard deviations for decomposition. We used multivariate linear mixed-effects models fitted through the *rma.mv()* function (METAFOR package) to calculate the pooled average SMD across all studies. In our multivariate linear mixed-effects models, we used a sampling error covariance matrix to account for the correlation between sampling errors within studies from which multiple effect sizes were extracted. To test whether decomposition was significantly affected by warming, we considered a pooled average effect size estimate to be significantly different from zero if the 95% CI around the mean did not include zero. Effects were considered significant at α=0.05.

To account for spatial autocorrelation between the study locations, we used a random effect consisting of longitude and latitude (in decimal degrees, with negative values for West and South) based on their great-circle distance (WGS84 ellipsoid method). In multivariate linear mixed-effects models, total heterogeneity (Q_T_) in effect sizes can be partitioned into heterogeneity explained by the model structure (Q_M_) and unexplained heterogeneity (Q_E_). We used the Test of Moderators (Q_M_ test) to determine how different factors, i.e., moderators, influence the effects of warming on decomposition (Koricheva *et al*. 2013).

We tested first for differences between the natural and the standardised plant litter dataset by using the data type (i.e., natural litter or standardised plant litter) as a moderator in the multivariate linear mixed-effects model. Because the effect of warming on decomposition of natural and standardised plant litter did not significantly differ (moderators’ test: Q_M_ (df = 2) = 2.7, p = 0.26), we combined the natural and standardised plant litter dataset in the following analyses and added the dataset type to the random-effects structure of our models to account for the different origins of our datasets (*random = ∼ LON + LAT | dataset type*).

To test the impact of macro-environment on the warming effect on decomposition, we first used multivariate linear mixed effects models (n=48) to explore whether the macro-environmental factors individually had a significant effect on the decomposition SMD (Table S6). However, as most environmental factors were confounded, we combined the macro-environmental factors to the underlying gradients using a Principal Component Analysis (PCA) on the scaled environmental variables using the R package FACTOMINER (v.2.4; Lê *et al*. 2008). We then used the four ‘macro-environmental classes’ created based on the origin of the PC1 and PC2 variables as a separation line, as moderator in the following multivariate linear mixed effects models to test whether the four environmental classes differed in their warming effect on decomposition. We used this factor ‘class’ as interacting moderator in the model to test for interactions in the macro-environment and the natural and standardised plant litter dataset.

We tested the impact of experimental induced changes in micro-environment, i.e., degree of warming and warming-induced changes in soil moisture and their interaction by testing them as moderators in the same multivariate linear mixed effects model (METAFOR package). We included the ‘macro-environmental class’ as interacting moderator to the model to test whether experimental induced changes in micro-environment differ between the four macro-environmental classes. To investigate whether experimental warming affected temperatures and soil moisture, we used a one-sample t-test to test whether the absolute difference between the warming treatment and the ambient control differed significantly from zero.

To test for differences in the warming effect between the different warming methods used in the different studies and experiments (Table S1, 2), we used ‘warming method’ as moderator in another multivariate linear mixed effects model. In this model, the macro-environmental class was not integrated because the warming methods were not evenly distributed across the four macro-environmental classes (e.g., more OTC studies in higher latitudes). To test for differences in the warming methods in their effect on micro-environment, we used linear mixed-effects models (R package LMERTEST, v. 3.1-3; Kuznetsova et al., 2017) to test the overall effect of the categorical independent variable ‘warming method’ on the continuous dependent variables ‘degree of warming’ and ‘warming-induced changes in soil moisture’, respectively. We used Tukey HSD post-hoc tests (R packages MULTCOMP, v. 1.4-19; Hothorn *et al*. 2008, and EMMEANS, v. 1.7.5; Lenth 2019) to check for significant differences between the warming methods in degree of warming and warming-induced changes in soil moisture, respectively. We further tested with a linear regression for correlations between warming-induced changes in soil moisture and the degree of warming.

To test for differences in litter quality, described as C:N ratio or ambient decomposability, between plant functional types in different macro-environments, we used linear mixed-effects models (R package LMERTEST) to test the overall effect of the categorical independent variables ‘plant functional type’ (including plant organ types) and ‘macro-environmental class’ and their interactions on the continuous dependent variables ‘C:N ratio’ or ‘ambient decomposability’ (different model for each). We then used Tukey HSD post-hoc tests (R packages MULTCOMP, v. 1.4-19; and EMMEANS, v. 1.7.5) to check for significant differences between the plant functional types and four macro-environmental classes in C:N ratio and ambient decomposability, respectively.

To test our hypothesis that lower litter quality is associated with a stronger positive warming effect on decomposition, we used multivariate linear mixed-effects models (METAFOR package) with the three applied proxies for litter quality ‘C:N ratio’, ‘ambient decomposability’ and ‘plant functional type’ (different model for each of the proxies) as moderators and ‘macro-environmental class’ again as an interactive factor.

In addition, we tested the site-specific drivers related to environmental conditions (absolute latitude and, altitude), experimental setup (duration of warming before the experiment, mesh size) as individual moderators fitting separate multivariate linear mixed-effects models (Table S5).

For each model, we tested the assumptions of normality and homogeneity of variance of the residuals. Whenever necessary, data were log-transformed (C:N ratio) or rank-transformed (warming-induced changes in soil moisture), with the latter case resulting in a non-parametric regression. Graphical displays were produced using the R packages GGPLOT2 (v. 3.3.6, Wickham *et al*. 2016) and ORCHARD (v.2.0, Nakagawa *et al*. 2021). We used R version 4.2.3 (R Core Team 2023) for all analyses.

#### Test for publication bias

Publication bias towards studies that report significant results is a common problem in meta-analysis (Sterne *et al*. 2000). We checked for publication bias in our literature data (on natural plant litter) by carrying out an Egger’s regression test for funnel plot asymmetry (using the *regtest* function, package METAFOR). The intercept of our regression model was 0.16 (CI95: - 0.07, 0.39). Because the 95% CI overlapped zero, no evidence for a publication bias was detected in our meta-analysis of decomposition responses to experimental warming (Egger *et al*. 1997; Sterne and Egger 2001).

## Results

### The effect of experimental warming on natural and standardised plant litter decomposition

The impact of experimental warming on plant litter decomposition was assessed across a range of temperatures of −1.6 to 7.5°C. On average, warming treatments significantly increased temperatures by 2.1 ± 0.1°C (mean ± SE, n=559; soil and air combined; t-test: t = 32.30, p < 0.001) and reduced soil moisture by 8.7 ± 0.9% (n=317; t-test: t = −9.23, p < 0.001. While the effect of warming on decomposition varied among studies, the overall standardised mean difference (SMD) did not significantly differ from zero (SMD = −0.08, p = 0.13 [CI95: −0.19, 0.02], *k*=637, Figure 2A). This pattern held for natural litter (SMD = −0.04, p = 0.58 [CI95: −0.20, 0.11], *k*=523; Figure 2B) and standardised plant litter (green tea: SMD = 0.14, p = 0.11 [CI95:-0.03, 0.32], *k*=57; rooibos tea: SMD = 0.08, p = 0.38 [CI95: 0.10, 0.26], *k*=57, Figure 2B). The effect of warming on decomposition did not significantly differ between natural and standardised plant litter datasets (moderators’ test: Q_M_ (df = 2) = 2.7, p = 0.26).

**Figure 2.**
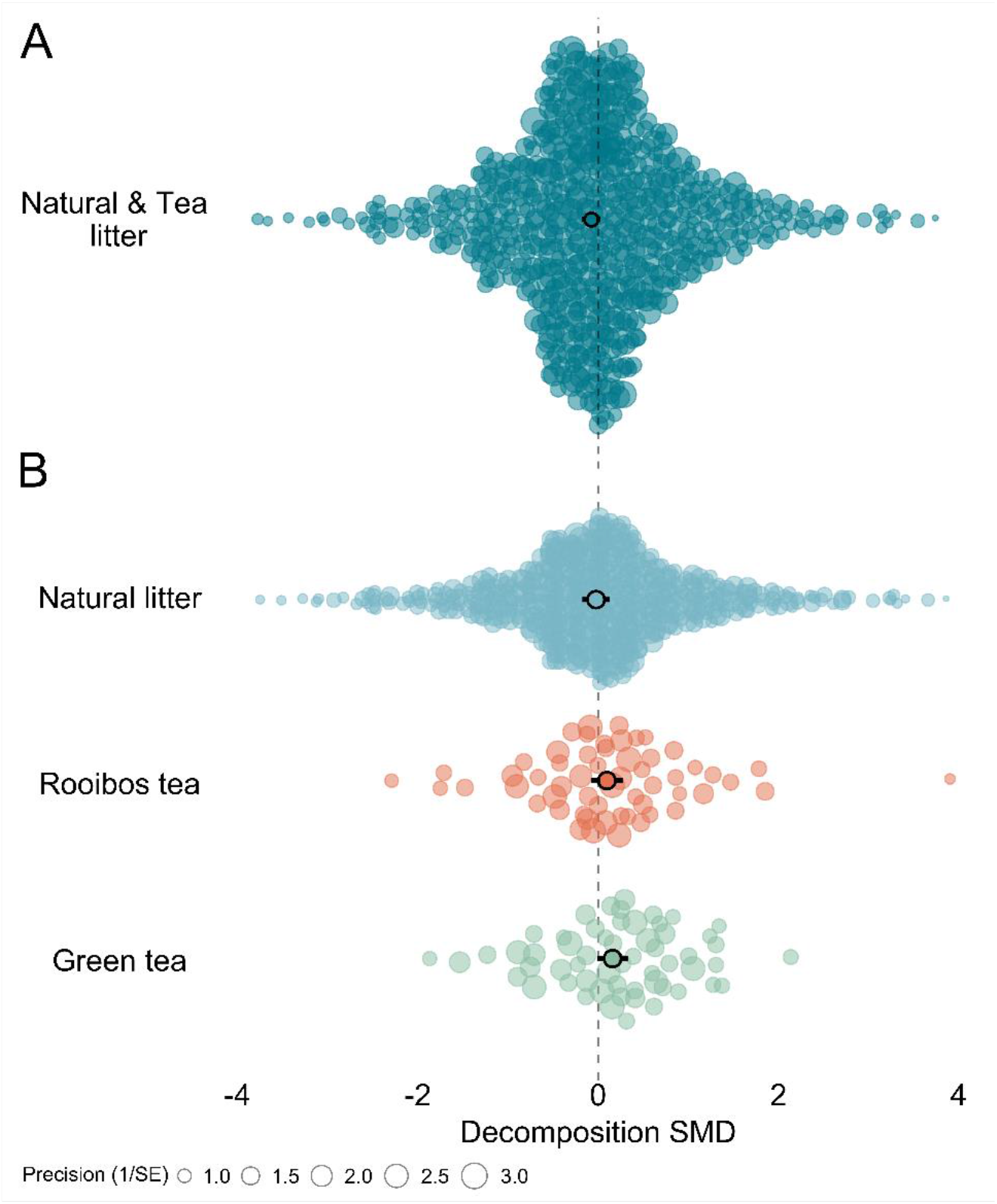
Effects of experimental warming on plant litter decomposition. The pooled average decomposition standardised mean difference (SMD, Hedges’ g; outlined circles) and 95% confidence intervals (black error bars) resulting from warming for (**A**) the whole dataset (i.e., natural and standardised plant litter; number of effect sizes *k*=637), and (**B**) separately for the natural litter (*k*=523), rooibos (*k*=57) and green tea (*k*=57). Each coloured dot is an individual effect size (non-outlined circles) with dot size representing its precision (the inverse of the standard error, larger points having greater influence on the model).

### The impact of macro-environment on the warming effect on decomposition

Macro-environmental factors individually tested had no influence on warming effects, except for specific factors (e.g., mean air temperature, aspect, deciduous tree and shrub cover; Table S6). However, the effect of warming on decomposition differed significantly across the four ‘macro-environmental classes’ identified by the PCA (moderators’ test: Q_M_ (df = 3) = 13.66, p = 0.003; Figure 3). Only, in the warm and dry class (high PC1 and low PC2 scores, Figure 3A), we observed a negative warming effect on decomposition (SMD = - 0.29, p = 0.01 [CI95: −0.51, −0.07], *k*=170; Figure 3B), driven primarily by natural litter (SMD = - 0.61, [CI95: −0.94; −0.28], *k*=150; Figure S3, Table S7). Despite the trend towards positive effects of warming on decomposition in the two cold classes, which comes with substantial variability, warming did not significantly affect decomposition in any of the other three macro-environmental classes and litter types (Figure 3B, Figure S3; Table S7).

**Figure 3.**
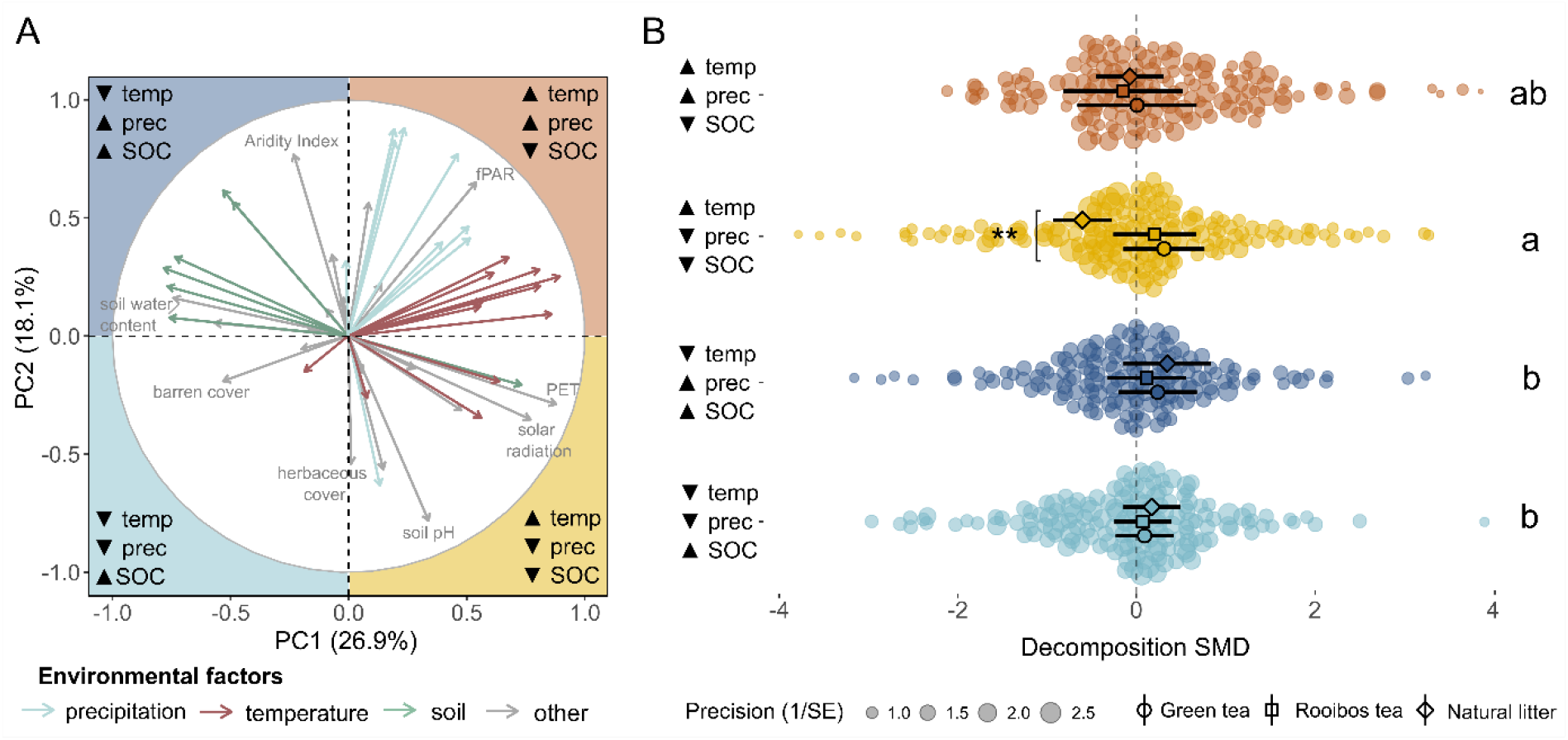
Impacts of macro-environment on decomposition responses to warming. (**A**) Principal component analysis (PCA) of the variation in macro-environmental factors in our dataset. The arrows are coloured according to the components, which are grouped into temperature, precipitation, soil and other factors (Table S3). The first two axes represent temperature and soil organic carbon-related variables (PC1), and precipitation (PC2). Full list of the macro-environmental factors, their scores on PC1 and PC2 and their mean in every class is presented in Table S3 and S4. Colours indicate the four macro-environmental classes distinguished by different combinations of high (▲) or low (▼) of temperature (temp), precipitation (prec) and soil organic carbon (SOC). (**B**) Pooled average decomposition SMD per macro-environmental class of natural litter (outlined diamonds), rooibos tea (outlines squares), and green tea (outlined circles) ±95%CI (error bars). Each coloured dot is an individual effect size (non-outlined circles) with dot size representing its precision (the inverse of the standard error, larger points having greater influence on the model). Asterisks indicate that the overall pooled average SMD is significantly different from zero (***p < 0.001), whereas different letters denote overall significant differences in the pooled average SMD across macro-environmental classes, averaged over data type.

### The impact of experimentally induced changes in micro-environment on decomposition and its interaction with macro-environment

The degree of warming correlated positively with the warming effect on decomposition: for every degree Celsius increase, the overall effect of warming on decomposition (SMD) increased by 0.18 (p < 0.001 [CI95: 0.10, 0.26], *k*=315; Figure S4). A significant increase in decomposition occurred with a degree of warming of 5.2 °C or more. However, the degree of warming’s impact on decomposition SMD depended on macro-environmental classes (moderators’ test: Q_M_ (df = 7) = 54.62, p < 0.001), with a more positive effect on decomposition in relatively warm and wet areas only (slope = 0.20 SMD/°C warming, p < 0.001 [CI95: 0.09, 0.31], *k*=315; Figure 4A).

**Figure 4.**
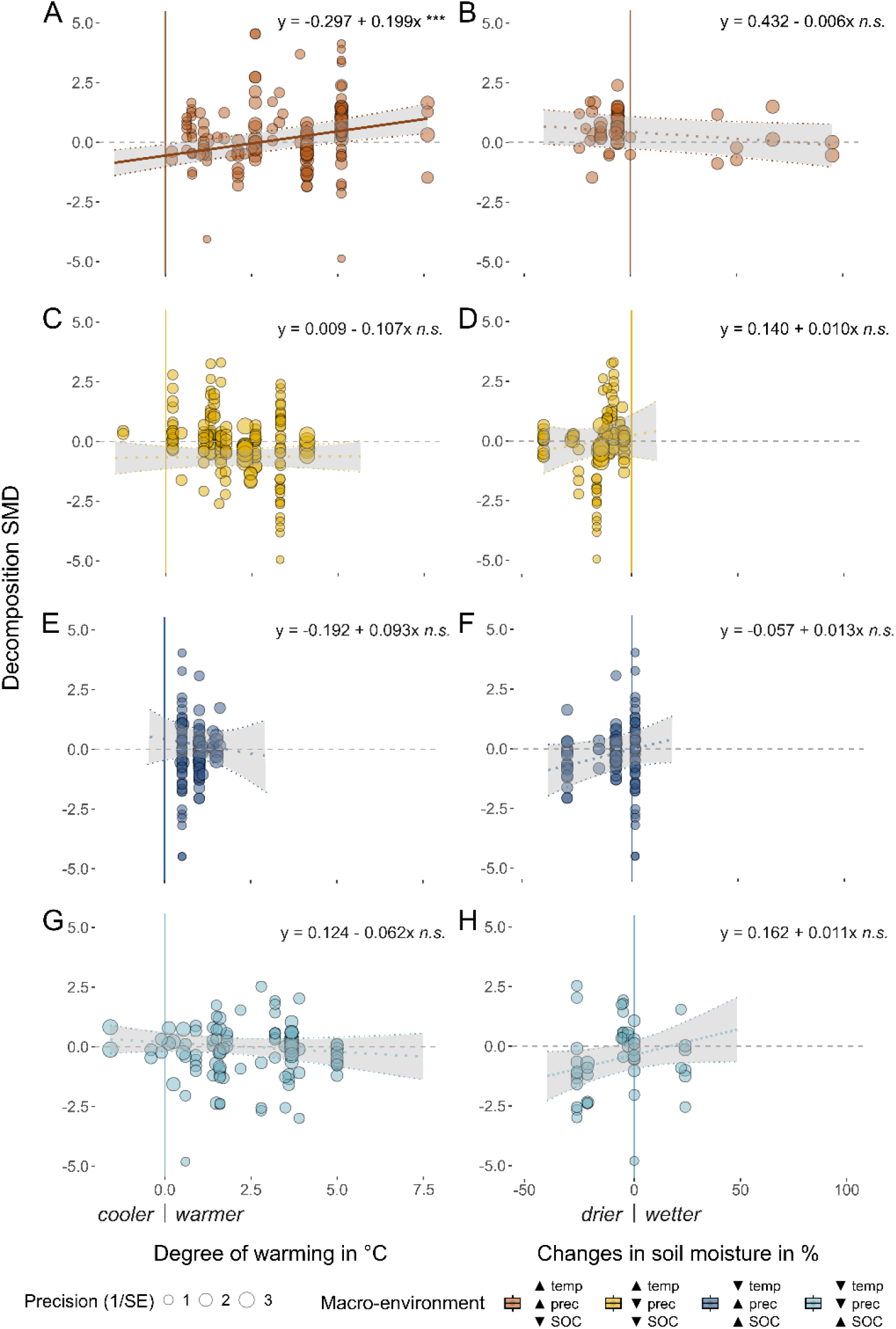
Relationships between the effect of warming on decomposition (SMD) and either (**A, C, E, G**) the degree of warming (i.e., absolute temperature difference between warmed and control plots) or (**B, D, F, H**) warming-induced changes in soil moisture (i.e., difference between warmed and control plots) separately for the four macro-environmental classes (different panels). Colours indicate the four macro-environmental classes distinguished by different combinations of high (▲) or low (▼) of temperature (temp), precipitation (prec) and soil organic carbon (SOC), consistent with Figure 3. Each coloured dot is an individual effect size (non-outlined circles) with dot size representing its precision (the inverse of the standard error, larger points having greater influence on the model). Solid lines indicate regression lines with shaded areas representing the 95%CI (***p < 0.001). Dashed lines indicate no significant relationship (n.s. = not significant).

In contrast, warming-induced changes in soil moisture had no impact on the warming effect on decomposition (p = 0.92 [CI95: −0.002, 0.002], *k*=315; Figure S4B-H) for any of the four macro-environmental classes (moderators’ test: Q_M_ (df = 7) = 13.63, p = 0.058). There was no significant interaction between degree of warming and changes in soil moisture in their impact of the warming effect on decomposition.

Warming methods significantly affected the warming effect on decomposition (moderators’ test: Q_M_ (df = 4) = 12.14, p = 0.016). Warming from heating cables significantly increased decomposition (SMD = 0.43, p = 0.010 [CI95: 0.10, 0.76], *k*=121) and was the most efficient warming (4.18 ± 0.1°C, n=121; Table 1), whichalso increased soil moisture (4.9 ± 4.1 %; n=48; Table 1). The warming effect of heating cables on decomposition differed significantly from the effect of OTCs on decomposition (Tukey HSD, p=0.006, Table 1), but was similar to the warming effect of infrared heaters or other warming methods on decomposition, none of which had a significant warming effect on decomposition (Table 1, Figure S5).

**Table 1.**
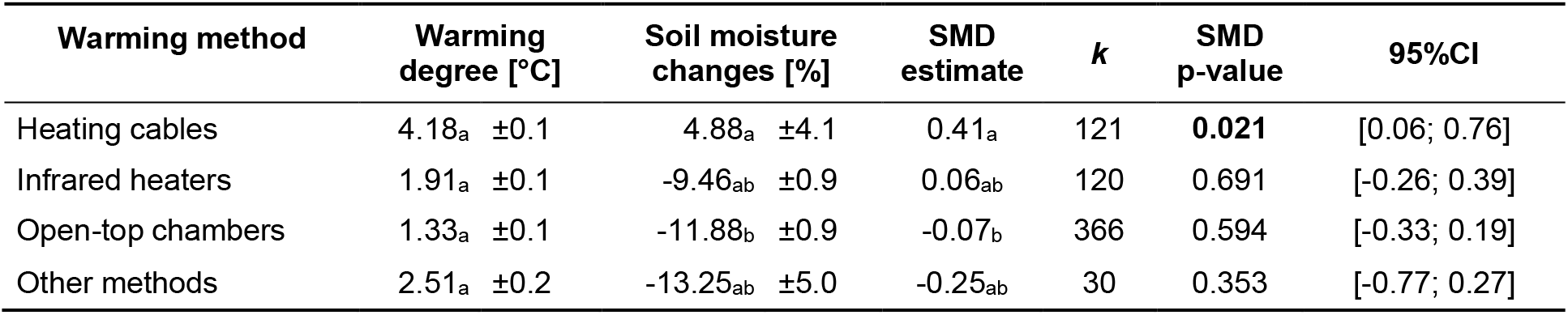
The impact of warming methods (i.e., Heating cables, Infrared heaters, Open-top chambers, other methods) on the degree of warming, the relative change in percent soil moisture (mean ± SE), and the effect of warming on decomposition (SMD). Different letters indicate significant differences in the degree of warming, the soil moisture changes, and the pooled average SMD across warming methods. Bold values indicate a significant effect of the warming method on SMD (p ≤ 0.05 or CI ≠ 0). Number of effect sizes (*k*), p-values for SMD estimates, and 95%-confidence interval are shown.

| Warming method | Warming degree [$^\circ\text{C}$] | Soil moisture changes [%] | SMD estimate | $k$ | SMD p-value | 95%CI |
| --- | --- | --- | --- | --- | --- | --- |
| Heating cables | 4.18<sub>a</sub> $\pm 0.1$ | 4.88<sub>a</sub> $\pm 4.1$ | 0.41<sub>a</sub> | 121 | **0.021** | [0.06; 0.76] |
| Infrared heaters | 1.91<sub>a</sub> $\pm 0.1$ | -9.46<sub>ab</sub> $\pm 0.9$ | 0.06<sub>ab</sub> | 120 | 0.691 | [-0.26; 0.39] |
| Open-top chambers | 1.33<sub>a</sub> $\pm 0.1$ | -11.88<sub>b</sub> $\pm 0.9$ | -0.07<sub>b</sub> | 366 | 0.594 | [-0.33; 0.19] |
| Other methods | 2.51<sub>a</sub> $\pm 0.2$ | -13.25<sub>ab</sub> $\pm 5.0$ | -0.25<sub>ab</sub> | 30 | 0.353 | [-0.77; 0.27] |

### The relationship between litter quality, ambient decomposability and the warming effect on decomposition

Plant functional types (including green and rooibos tea) and plant organ types differed significantly in their C:N ratio (ANOVA: F(9, 72) = 417.9, p < 0.001, n = 72; Figure S6A). While the C:N ratio was not significantly related to the warming effect on decomposition (slope=-0.002 SMD/C:N ratio, p = 0.27 [CI: −0.006, 0.002], k=428), plant functional types and organ types differed in their warming effect on decomposition (moderators’ test: Q_M_ (df = 8) = 47.92, p < 0.001). Warming increased decomposition of graminoid roots (SMD = 0.55, p<0.001 [CI:0.27, 0.84], k=49) and decreased decomposition of graminoid shoots and leaves (SMD = - 0.25, p=0.010 [CI: −0.43, −0.06], *k*=151). Woody species showed no significant warming effect on broadleaves or roots but warming significantly decreased decomposition of woody needle litter (SMD = −0.44, p=0.021 [CI: −0.82, −0.07], *k*=20; Figure S6B, Table S8). Although data were not available for all plant functional types and plant organ types across all four macro-environmental classes, the available data indicates that the macro-environment determined how decomposition of different plant functional types responded to warming (moderators’ test: Q_M_ (df = 28) = 138.82, p < 0.001).

Data on woody roots were available only for the warm and wet class where woody roots’ decomposition increased with warming (SMD = 1.08, p=0.040 [CI95: 0.05, 2.11], *k*=5, Figure 5A). In the warm and dry class, only moss- and lichen litter decomposition significantly increased with warming (SMD = 1.10, p = 0.026 [CI95: 0.13, 2.07], *k*=12, Figure 5B). In the cold and wet class, warming decreased decomposition of woody broadleaf litter (SMD = −0.20, p = 0.010 [CI95: −0.35, −0.05], *k*=66) and green tea (SMD = 0.34, p = 0.016 [CI95: 0.06, 0.61], *k*=15, Figure 5C). Lastly, in the cold and dry class, only graminoids roots’ decomposition significantly increased with warming (SMD = 0.95, p=0.049 [CI95: 0.004, 1.89], *k*=7, Figure 5D).

**Figure 5.**
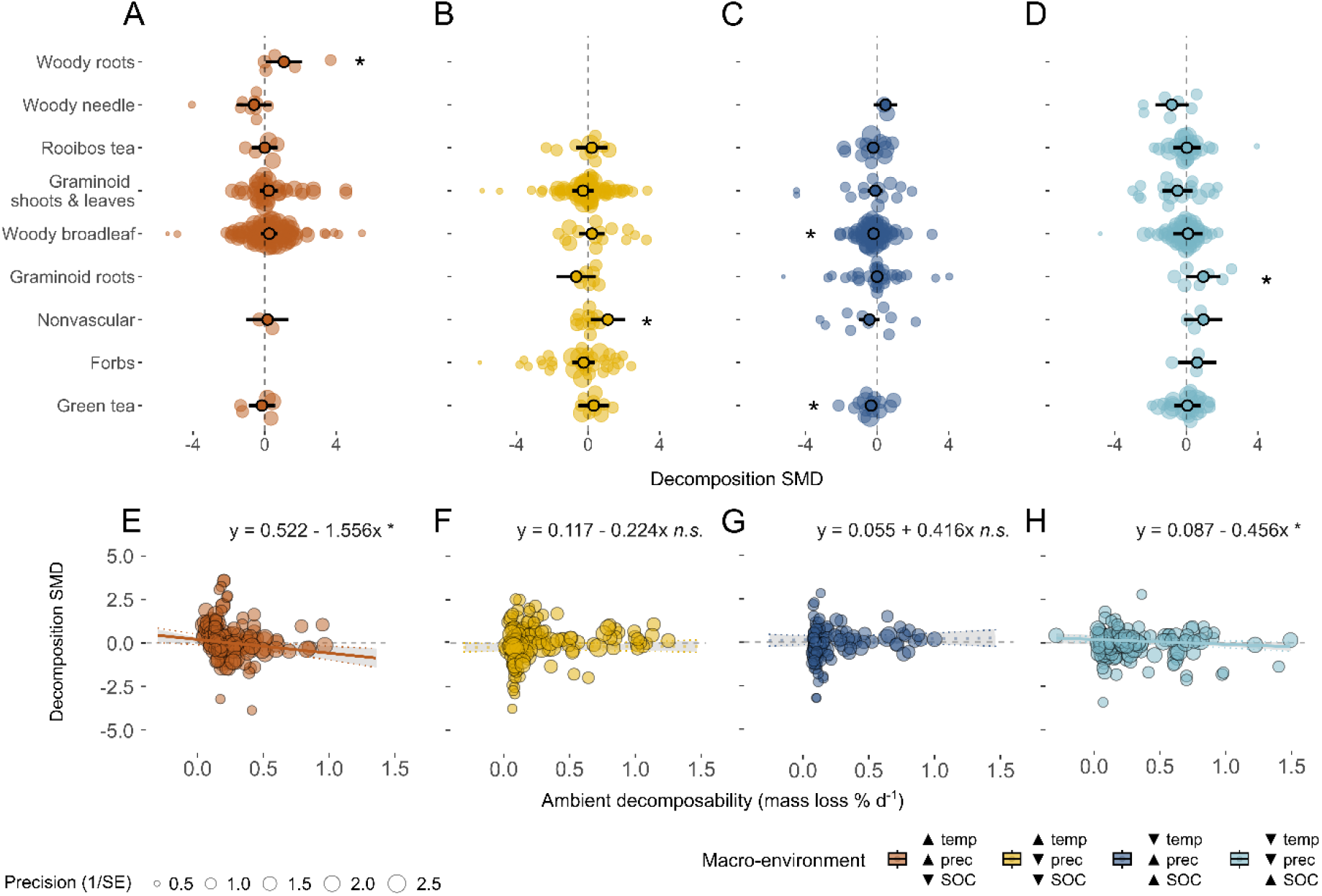
Relationship between litter quality, ambient decomposability and the warming effect on decomposition across the four macro-environmental classes. (**A, B, C, D**) The pooled average decomposition standardized mean difference (SMD, Hedges’ g, black outlined circles) and 95% confidence intervals (black error bars) for different plant functional types (in cases data was available; see methods) in each of the four macro-environmental classes. (**E, F, G, H**) Relationship between ambient decomposability (ambient mass loss rate in % d^-1^) and the warming effect on decomposition for each of the four macro-environmental classes (see also Figure S7). Colours indicate the four macro-environmental classes of temperature (temp), precipitation (prec) and soil organic carbon (SOC) that are either high (▲) or low (▼), consistent with Figure 3. Each coloured dot is an individual effect size (non-outlined circles) with dot size representing its precision (the inverse of the standard error, larger points having greater influence on the model). Solid lines indicate regression lines with shaded areas representing the 95%CI. Asterisks indicate that the overall pooled average SMD is significantly different from zero (*p < 0.05). Dashed lines indicate no significant relationship (n.s. = not significant).

Overall, the effect of warming on decomposition was more positive for litter that decomposed slower under ambient conditions (i.e., lower decomposability) (moderators’ test: Q_M_ (df = 1) = 8.58, p = 0.003). Whereas ambient decomposability was on average equal in the four macro-environmental classes (ANOVA: F(3, 124) = 1.21, p = 0.31), its importance as a driver for the warming effect was only significant in the warm and wet macro-environmental class (slope = −0.31 SMD/ decomposability, p = 0.029 [CI95: −0.58, −0.03], *k*=158; Figure 5E) and in the cold and dry class (slope = 0.28 SMD/ decomposability, p = 0.017 [CI95: −0.69, - 0.14], *k*=151; Figure 5H).

## Discussion

Across 109 datasets and 637 paired observations of plant litter decomposition under warmed vs. ambient conditions across all litter types on global scale, we found that the overall effect of warming only increased decomposition when warming exceeded 5.2°C and moisture is not limited. This estimated threshold is beyond the global warming predicted by the end of the century (1.4 to 4.4°C; IPCC 2021). As expected, the macro-environmental region is a key determinant of the effect of warming on decomposition and especially in warm, moisture-limited systems decomposition decreased under warming. The quality of the litter was an important moderator of the warming effect, especially in warm and wet as well as cold and dry conditions. The decomposition of slowly degradable material increased with warming, while the decomposition of easily degradable material decreased with warming.

### Context-dependencies of the warming effect on decomposition

Our findings confirm that effects of climate change on decomposition may depend strongly on future warming magnitude and confounded impact on moisture (Aerts 2006). Consequently, we found that the prevailing macro-environmental conditions have a critical influence on whether warming leads to an increase, a decrease or no change in litter decomposition. As hypothesized, warming reduced decomposition at warmer sites with limited water availability due to low precipitation and high evapotranspiration (Sierra *et al*. 2017).

Macro-environmental conditions in warmer regions such as temperate and subtropical areas, generally favor decomposition processes (Meentemeyer 1978; Powers *et al*. 2009). Hence, in warm ecosystems, the potential for accelerating decomposer activity through experimental warming may be relatively limited (Bradford 2013; Crowther and Bradford 2013). Instead, the role of soil moisture becomes a potentially more important limiting factor (Aerts 2006). Accordingly, our observation that warming led to decreased decomposition in warm and dry areas, but not in warm and wet areas, aligns with both our expectations and previous studies where warming amplified drought in dry macro-environments but not in wet ones (Wu *et al*. 2011; Thakur *et al*. 2018; Schimel 2018). It is noteworthy that warm and dry systems exhibit the lowest soil organic carbon content (Table S4), indicating limited carbon storage potential in these warm and dry sites. This implies that while warming decreases decomposition in warm and dry systems, the effectiveness of carbon storage is likely compromised due to warmer temperatures and dry conditions (Yi *et al*. 2014; Hartley *et al*. 2021). While previous studies have reported clear interactions between increasing temperature and moisture (Aerts 2006; Thomas *et al*. 2023), our research shows that these interactions manifest differently in various macro-environments. We emphasize how the predominant macro-environment interacts with site-specific factors, such as the degree of warming, a significant moderator only in the warm and wet class but not in other classes.

Contrary to our expectations, we did not observe significant effects of warming on decomposition in studies conducted at colder sites, where temperature, rather than water availability, appears to be the main constraint. Despite a tendency for positive effects in the cold macro-environment class (wet and dry), it is surprising that a significant positive effect of warming in colder systems was not found. Given that we found some evidence for higher temperature sensitivity of recalcitrant litter (Biasi *et al*. 2005; Davidson and Janssens 2006; Conant, Steinweg, *et al*. 2008), and that cold systems (i.e., high latitude and alpine regions with tundra and boreal forests; Figure S2B) are dominated by more recalcitrant plant functional types (Dorrepaal *et al*. 2005), as well as having a greater proportion of plant material below ground (Mokany *et al*. 2006; Poorter *et al*. 2012; Iversen *et al*. 2015; Wang *et al*. 2016), we expected a significant positive effect of warming in colder systems. In addition, we might not have been able to detect any significant effect in these cold macro-environments due to passive warming methods (e.g., OTCs) prevailing there (Aronson and McNulty 2009). These warming methods do typically not achieve the degree of warming of 5.2°C that we estimated to be necessary to find significant positive effects on decomposition. However, warming in high latitude and high-altitude systems has outpaced the global average, with the Arctic warming nearly four times faster over the last four decades (Tingley and Huybers 2013; Rantanen *et al*. 2022). This suggests that our predicted temperature threshold could become relevant for these systems and that ongoing future warming may potentially accelerate decomposition in these environments with exceptionally high carbon storage potential.

### Litter quality and ambient decomposability as regulators of the warming effect on decomposition

We found that litter material that was harder to decompose under ambient conditions, decomposed faster under warming compared to litter material that was easy to decompose under ambient conditions, which was significant under warm and wet as well as cold and dry conditions. As lower decomposability is frequently associated with lower quality litter (Cornelissen *et al*. 2004; Freschet *et al*. 2012), this supports our hypothesis that lower quality litter is more sensitive to warming than high quality litter (Bosatta and Ågren 1999; Fierer *et al*. 2005; Conant, Drijber, *et al*. 2008; Suseela *et al*. 2013). Therefore, we found indications that warming could disproportionately accelerate the decomposition of litter with a low decomposability (Biasi *et al*. 2005; Davidson and Janssens 2006; Conant, Steinweg, *et al*. 2008). Of concern is that these lower quality litter types with more recalcitrant material are more important for carbon input into the soil (Parton *et al*. 1987; Davidson and Janssens 2006). An increase in their decomposition, provided that the carbon uptake of plants does not surpass the decomposition rate, may result in a decline in net inputs to soil organic matter storage (Hicks Pries *et al*. 2015). This potential decrease in carbon sequestration rates could contribute to the acceleration of climate change by releasing more carbon into the atmosphere as CO_2_ (Jenkinson *et al*. 1991; Kirschbaum 1995, 2000; Cox *et al*. 2000; Fenner and Freeman 2011). Surprisingly, the warming effect on decomposition appears to be unrelated to a classic measure of litter quality (C:N ratio). We observed the strongest correlation of the warming effect on decomposition with ambient decomposability, which integrates both litter quality and environmental conditions (Cornelissen *et al*. 2004; Freschet *et al*. 2012). While the quality of the litter material (C:N ratio) may not strongly drive decomposition under experimental warming, our study emphasizes the importance of considering the interaction between litter quality and environmental conditions for understanding decomposition dynamics in response to climate change (Joly *et al*. 2023). The varying effects of warming on the decomposition of different plant functional litter types and plant organs suggest that the specific structural composition of plant species or functional groups within a plant community significantly influences warming responses.

Specifically, we found that, warming led to decreased decomposition for shoots and leaves, while it increased decomposition for roots (Figure S6B). This contrasting response might be due to inherent structural traits in roots, such as lignin, carbon, and dry matter content, making them more resistant to decomposition (Freschet *et al*. 2012; Xia *et al*. 2015), consequently, more temperature-sensitive and responsive to warming (Bosatta and Ågren 1999; Fierer *et al*. 2005; Conant, Drijber, *et al*. 2008; Suseela *et al*. 2013). In addition, the distinct responses of shoot/leave and root decomposition to warming might be partly attributed to their incubation position in the soil or on the soil surface (Blok *et al*. 2018, Table S8). The specific environmental conditions in which above and below ground decomposition occurs likely influence the response to warming, with soil serving as a buffer against extreme conditions (Wang *et al*. 2009; Fanin *et al*. 2020). Drier soil surface conditions likely contributed to the negative impact of warming on leaf and shoot decomposition (Blok *et al*. 2018), whereas wetter soil conditions enhanced decomposition under warming, irrespective of the plant organ type (Hicks Pries *et al*. 2013). The distinct impact of warming on roots and shoots/leaves is unlikely to be caused by differences in litter quality since the C:N ratio did not differ between roots and shoots/leaves. This opposite response of plant organ types was exclusive to graminoids and not observed in woody species, indicating a yet undiscovered potential interaction between warming and plant functional type. This urges further investigation especially for accurate assessments of carbon and nutrient budgets in a warming climate since root production and turnover accounts for 20-80 % of the global annual net primary productivity (Jackson *et al*. 1997; McCormack *et al*. 2015). This knowledge gap currently posing a challenge for carbon cycle modelling especially in ecosystems with a significant portion of biomass located below ground.

### Limitations of specific warming methods

By including a range of warming methods, we were able to overcome some of the limitations or confounding effects associated with specific warming methods. Heating cables were the most effective warming methods that did not affect soil moisture and led to a significant increase in decomposition. Heating cables dominated in the warm and wet macro-environmental class, which was the only class where the degree of warming was a significant moderator and exceeded the threshold of 5.2°C that we estimated was necessary to find significant positive effects on decomposition. However, by warming soils rather than air, heating cables may provide less realistic warming conditions. Infrared heaters rather replicate natural warming conditions (Aronson and McNulty 2009), but had no effect on decomposition as their warming capacity was relatively small. The non-significant effect of passive warming by OTCs on litter decomposition might be explained by confounding factors, such as reduced soil temperatures due to shade or increased radiation absorption in OTCs, resulting in less effective warming, but increased drought effects on the soil (Marion *et al*. 1997; Aerts 2006; Shu *et al*. 2019; Hollister *et al*. 2023). In our dataset, OTCs were the most common warming method in cold ecosystems, but were associated with the largest decrease in soil moisture (- 11.88 ± 0.9 %) and a relatively small but realistic warming (+1.33 ± 0.1°C; Table 1). By combining studies of different warming methods, we were able to demonstrate the context-dependency of the warming effect with the microenvironment. We found a significant warming effect on decomposition only for warming methods (i.e., heating cables) with a high degree of warming that did not decrease soil moisture, but actually increased it (Table 1).

Our data set covers large parts of the world and most biomes (Figure 1), but we lack data on tropical and temperate rainforests in our dataset. Either because experimental warming *in-situ* studies in these biomes are not conducted or did not match our criteria of the literature survey. Hence, in our study, the impact of warming on rainforest decomposition remains uncertain and we suggest it should be a priority for future research.

### Global implications

This global meta-analysis integrates all available data from *in situ* experimental warming studies on decomposition across terrestrial ecosystems worldwide. The global approach enabled us the exploration of context-dependencies between warming effects and various environmental factors, such as moisture and plant functional types, which is limited for single studies performed on mainly regional scales. We show that predictions of the ongoing impacts of climate change on key ecosystem processes should take into account the context-dependencies of macro-environment and litter quality.

In particular, the effects of warming on below-ground decomposition need to be further explored, as our results indicate that there may be an important, yet poorly investigated, context-dependent interaction between below-ground litter decomposition (i.e. roots) and warming that is distinct from the warming effects of above-ground litter (i.e. shoots). This poses challenges for carbon cycle modeling, especially in regions like tundra, cold deserts, and temperate grasslands, where up to 80 % of the plant biomass is located below ground (Mokany *et al*. 2006; Poorter *et al*. 2012; Iversen *et al*. 2015; Wang *et al*. 2016). Furthermore, many biomes are currently undergoing rapid vegetation changes. The lack of data, particularly for woody species makes it difficult to predict how nutrient and carbon cycling will change with warming for tundra, alpine systems and savannahs, where increases in shrub cover with relatively recalcitrant litter have been observed (Harte *et al*. 1995; Myers-Smith *et al*. 2011; Elmendorf *et al*. 2012; Pearson *et al*. 2013; García Criado *et al*. 2020). Our results suggest that these changes in vegetation may lead to higher carbon release and lower input of soil organic matter, reducing storage potential through increased decomposition under warming of lower quality plant litter.

Based on our finding, litter decomposition is only likely to increase significantly under more extreme warming scenarios in the range of 3.3°C to 5.7°C (SSP5-8.5, IPCC 2021). This becomes increasingly relevant, considering that we faced the warmest years on record for the globe as a whole in the last decade. It further has direct relevance for high-latitude systems that experience the most rapid warming, with an average temperature increase of 0.65 ± 0.09°C per decade (from 1979 to 2022) according to ERA5 (ECMWF Reanalysis v5, European Centre for Medium-Range Weather Forecasts).

Furthermore, the significant increases in droughts experienced in Europe, the Mediterranean region and large parts of Asia signify that additional warming leading to drier soils might increasingly become a limiting factor (NOAA National Centers for Environmental Information). Based on our findings, the anticipated rise in the intensity and frequency of drought events, driven by climate change, may result in reduced decomposition rates in this warm and dry regions in the future. Depending on how drought will affect primary productivity, this could lead to reduced soil carbon emissions from the soil. However, our findings suggest that this effect is more prominent in ecosystems with inherently lower initial carbon storage potential, which might indicate that warming effects on decomposition in these warm and dry systems may play a minor role for worldwide carbon budgets.

Certainly, the net carbon balance of a system is as well determined by carbon uptake, yet decomposition plays a pivotal role in the carbon budget. This study improves our understanding of the context-dependence of warming sensitivity, contributing to more accurate predictions of climate change impacts on decomposition as a key ecosystem process.

## ACKNOWLEDGEMENTS

We would like to acknowledge the numerous students and field assistants including Bin Xu, Courtney Campbell, Sasha van Stavel, Jordanna Branham and Golnoush Fard, involved in the collection of tea measurements and like to thank station managers for their help and access to field sites. We thank Melanie Bird for her assistance in the laboratory and Albin Bjärhall for his support in extracting raw data from the published literature. Recognizing the importance of Indigenous lands, we acknowledge that parts of our fieldwork were conducted on territories historically and presently belonging to Indigenous peoples. We express our respect and gratitude to these communities. Special thanks are extended to the residents of Utqiagvik and Atqasuk, Alaska, for their cooperation and understanding during our research activities in the Arctic region. This research would not have been possible without the collective efforts and support of these individuals and communities.

## FUNDING

For SS and JMS funding was received from Formas (Grant No: 2021-02449). JMS also acknowledges support from the Swedish Research Council VR (Grant No: 2014-04270).

Support for ED was provided by the Swedish Research Council VR (Grant No: 2018-04004). The research conducted by MS and TM was funded by Alberta Innovates Technology Futures. Additionally, MS acknowledges support from an NSERC Canada Research Chair (CRC-2019-00299).

EA was an International Research Fellow of the Japan Society for the Promotion of Science (JSPS) with ID No: P17102.

Funding for IA and VV was provided by the Research Council of Norway under the KLIMAFORSK program (Grant No: 244525).

MPB acknowledges support from the Governor of Svalbard (Svalbard Environmental Protection Fund, Grant Project No: 15/128), the Research Council of Norway (Arctic Field Grant, Project No: 269957), and the National Science Foundation (Grant Project No: ANS-2113641).

R Alonso acknowledges funding from the Framework on atmospheric pollution and persistent organic pollutants between DGCEA and CIEMAT (ACTUA-MITERD).

NF was funded by a grant for the organization of a new laboratory for young researchers at Yugra State University as part of the implementation of the National Project "Science and Universities."

RDH acknowledges support from the US National Science Foundation (Grant No: 1836839). IJS was funded by the University of Iceland Research Fund for the years 2016 and 2017.

QL acknowledges the CAS International partnership project (Grant No: 131323KYSB20210004).

IM acknowledges funding from the UK Natural Environment Research Council for the ShrubTundra Project (Grant No: NE/M016323/1).

YY was supported by the Sichuan Provincial Science and Technology Plan Project (Grant No: 2022ZHYZ0005).

## Statement of authorship

JMS conceived the idea and SS, JMS, and ED designed the study. JMS, ED, MPB, VV, ElR, and MS gave extensive feedback on the analyses and manuscript. RA, EA, JA, IA, KB, SB, MPB, MC, AC, CTC, KC, R Alonso, NF, WG, IAH, AH, RDH, ISJ, EHK, KK, QL, JOL, AM, TM, IM, AP, MS, S Suzuki, TK, MtB, HT, FV, VV, SV, YY, J Asplund, and ElR collected the data on standardised plant litter decomposition from open-top-chamber warming experiments. Jvd and TWC provided the map-based environmental data. All authors excluding SS, ED, and JS contributed data. SS and JMS assembled the data for meta-analysis and meta-regression. SS analysed the data with feedback from JMS and ED. SS designed the figures and tables, and wrote the manuscript.

## Data availability statement

The data that support the findings of this study will be made openly available on Dryad at https://doi.org/10.5061/dryad.p5hqbzkw5 once accepted for publication.

## References

Aerts R. 1997. Climate, Leaf Litter Chemistry and Leaf Litter Decomposition in Terrestrial Ecosystems: A Triangular Relationship. Oikos 79: 439.

Aerts R. 2006. The freezer defrosting: global warming and litter decomposition rates in cold biomes: *Global warming and litter decomposition*. Journal of Ecology 94: 713–724.

Anderegg WR, Diffenbaugh NS. 2015. Observed and projected climate trends and hotspots across the National Ecological Observatory Network regions. Frontiers in Ecology and the Environment 13: 547–552.

Aronson EL, McNulty SG. 2009. Appropriate experimental ecosystem warming methods by ecosystem, objective, and practicality. Agricultural and Forest Meteorology 149: 1791–1799.

Bardgett RD, Freeman C, Ostle NJ. 2008. Microbial contributions to climate change through carbon cycle feedbacks. The ISME Journal 2: 805–814.

Bardgett RD, Manning P, Morriën E, De Vries FT. 2013. Hierarchical responses of plant-soil interactions to climate change: consequences for the global carbon cycle (W Van Der Putten, Ed.). Journal of Ecology 101: 334–343.

Berbeco MR, Melillo JM, Orians CM. 2012. Soil warming accelerates decomposition of fine woody debris. PLANT AND SOIL 356: 405–417.

Biasi C, Rusalimova O, Meyer H, et al. 2005. Temperature-dependent shift from labile to recalcitrant carbon sources of arctic heterotrophs. Rapid Communications in Mass Spectrometry 19: 1401–1408.

Björnsdóttir K, Barrio IC, Jónsdóttir IS. 2022. Long-term warming manipulations reveal complex decomposition responses across different tundra vegetation types. Arctic Science 8: 979–991.

Blok D, Faucherre S, Banyasz I, Rinnan R, Michelsen A, Elberling B. 2018. Contrasting above- and belowground organic matter decomposition and carbon and nitrogen dynamics in response to warming in High Arctic tundra. GLOBAL CHANGE BIOLOGY 24: 2660–2672.

Bosatta E, Ågren GI. 1999. Soil organic matter quality interpreted thermodynamically. Soil Biology and Biochemistry 31: 1889–1891.

Bradford M. 2013. Thermal adaptation of decomposer communities in warming soils. Frontiers in Microbiology 4.

Chapin FS, Bret-Harte M, Hobbie S, Zhong H. 1996. Plant functional types as predictors of transient responses of arctic vegetation to global change. Journal of Vegetation Science 7.

Chen J, Luo Y, Xia J, et al. 2015. Stronger warming effects on microbial abundances in colder regions. Scientific Reports 5: 18032.

Cheng X, Luo Y, Su B, et al. 2010. Experimental warming and clipping altered litter carbon and nitrogen dynamics in a tallgrass prairie. AGRICULTURE ECOSYSTEMS & ENVIRONMENT 138: 206–213.

Christiansen C, Haugwitz M, Prieme A, et al. 2017. Enhanced summer warming reduces fungal decomposer diversity and litter mass loss more strongly in dry than in wet tundra. GLOBAL CHANGE BIOLOGY 23: 406–420.

Conant RT, Drijber RA, Haddix ML, et al. 2008. Sensitivity of organic matter decomposition to warming varies with its quality. Global Change Biology 14: 868–877.

Conant RT, Ryan MG, Ågren GI, et al. 2011. Temperature and soil organic matter decomposition rates – synthesis of current knowledge and a way forward. Global Change Biology 17: 3392–3404.

Conant RT, Steinweg JM, Haddix ML, Paul EA, Plante AF, Six J. 2008. Experimental warming shows that decomposition temperature sensitivity increases with soil organic matter recalcitrance. Ecology 89: 2384–2391.

Cornelissen JHC, Quested HM, Gwynn-Jones D, et al. 2004. Leaf digestibility and litter decomposability are related in a wide range of subarctic plant species and types. Functional Ecology 18: 779–786.

Coûteaux M-M, Bottner P, Berg B. 1995. Litter decomposition, climate and liter quality. Trends in Ecology & Evolution 10: 63–66.

Cox PM, Betts RA, Jones CD, Spall SA, Totterdell IJ. 2000. Acceleration of global warming due to carbon-cycle feedbacks in a coupled climate model. Nature 408: 184–187.

Crowther TW, Bradford MA. 2013. Thermal acclimation in widespread heterotrophic soil microbes. Ecology Letters 16: 469–477.

Davidson EA, Janssens IA. 2006. Temperature sensitivity of soil carbon decomposition and feedbacks to climate change. Nature 440: 165–173.

Davidson EA, Trumbore SE, Amundson R. 2000. Soil warming and organic carbon content. Nature 408: 789–790.

Djukic I, Kepfer-Rojas S, Schmidt IK, et al. 2018. Early stage litter decomposition across biomes. Science of The Total Environment 628–629: 1369–1394.

Dorrepaal E, Cornelissen JHC, Aerts R, Wallén B, Van Logtestijn RSP. 2005. Are growth forms consistent predictors of leaf litter quality and decomposability across peatlands along a latitudinal gradient?: Plant growth forms and litter quality. Journal of Ecology 93: 817–828.

Egger M, Smith GD, Schneider M, Minder C. 1997. Bias in meta-analysis detected by a simple, graphical test. BMJ 315: 629–634.

Elmendorf SC, Henry GHR, Hollister RD, et al. 2012. Plot-scale evidence of tundra vegetation change and links to recent summer warming. Nature Climate Change 2: 453–457.

Fanin N, Bezaud S, Sarneel JM, Cecchini S, Nicolas M, Augusto L. 2020. Relative Importance of Climate, Soil and Plant Functional Traits During the Early Decomposition Stage of Standardized Litter. Ecosystems 23: 1004–1018.

Fenner N, Freeman C. 2011. Drought-induced carbon loss in peatlands. Nature Geoscience 4: 895– 900.

Fierer N, Craine JM, McLauchlan K, Schimel JP. 2005. Litter Quality and the Temperature Sensitivity of Decomposition. Ecology 86: 320–326.

Fontaine S, Bardoux G, Abbadie L, Mariotti A. 2004. Carbon input to soil may decrease soil carbon content. Ecology Letters 7: 314–320.

Freschet GT, Aerts R, Cornelissen JHC. 2012. A plant economics spectrum of litter decomposability. Functional Ecology 26: 56–65.

García Criado M, Myers-Smith IH, Bjorkman AD, Lehmann CER, Stevens N. 2020. Woody plant encroachment intensifies under climate change across tundra and savanna biomes. Global Ecology and Biogeography 29: 925–943.

Gong S, Guo R, Zhang T, Guo J. 2015. Warming and Nitrogen Addition Increase Litter Decomposition in a Temperate Meadow Ecosystem. PLOS ONE 10.

Harte J, Torn MS, Fang-Ru Chou Chang, et al. 1995. Global Warming and Soil Microclimate: Results from a Meadow-Warming Experiment. Ecological Applications 5: 132–150.

Hartley IP, Hill TC, Chadburn SE, Hugelius G. 2021. Temperature effects on carbon storage are controlled by soil stabilisation capacities. Nature Communications 12: 6713.

Hedges LV. 1981. Distribution Theory for Glass’s Estimator of Effect size and Related Estimators. Journal of Educational Statistics 6: 107–128.

Henry HAL, Moise ERD. 2015. Grass litter responses to warming and N addition: temporal variation in the contributions of litter quality and environmental effects to decomposition. PLANT AND SOIL 389: 35–43.

Hicks Pries CE, Logtestijn RSP, Schuur EAG, et al. 2015. Decadal warming causes a consistent and persistent shift from heterotrophic to autotrophic respiration in contrasting permafrost ecosystems. Global Change Biology 21: 4508–4519.

Hicks Pries CE, Schuur E a. G, Vogel JG, Natali SM. 2013. Moisture drives surface decomposition in thawing tundra. Journal of Geophysical Research: Biogeosciences 118: 1133–1143.

Hollister RD, Elphinstone C, Henry GHR, et al. 2023. A review of open top chamber (OTC) performance across the ITEX Network. Arctic Science 9: 331–344.

Hong J, Lu X, Ma X, Wang X. 2021. Five-year study on the effects of warming and plant litter quality on litter decomposition rate in a Tibetan alpine grassland. SCIENCE OF THE TOTAL ENVIRONMENT 750.

Hothorn T, Bretz F, Westfall P. 2008. Simultaneous inference in general parametric models. Biometrical Journal 50: 346–363.

Iversen CM, Sloan VL, Sullivan PF, et al. 2015. The unseen iceberg: plant roots in arctic tundra. New Phytologist 205: 34–58.

Jackson RB, Mooney HA, Schulze E-D. 1997. A global budget for fine root biomass, surface area, and nutrient contents. Proceedings of the National Academy of Sciences 94: 7362–7366.

Jenkinson DS, Adams DE, Wild A. 1991. Model estimates of CO2 emissions from soil in response to global warming. Nature 351: 304–306.

Joly F-X, Scherer-Lorenzen M, Hättenschwiler S. 2023. Resolving the intricate role of climate in litter decomposition. Nature Ecology & Evolution 7: 214–223.

Keuskamp JA, Dingemans BJJ, Lehtinen T, Sarneel JM, Hefting MM. 2013. Tea Bag Index: a novel approach to collect uniform decomposition data across ecosystems (H Muller-Landau, Ed.). Methods in Ecology and Evolution 4: 1070–1075.

Kirchman DL. 2018. Degradation of organic matter. Oxford University Press.

Kirschbaum MUF. 1995. The temperature dependence of soil organic matter decomposition, and the effect of global warming on soil organic C storage. Soil Biology and Biochemistry 27: 753–760.

Kirschbaum MUF. 2000. Will changes in soil organic carbon act as a positive or negative feedback on global warming? Biogeochemistry 48: 21–51.

Kirschbaum MUF. 2004. Soil respiration under prolonged soil warming: are rate reductions caused by acclimation or substrate loss? Global Change Biology 10: 1870–1877.

Knorr W, Prentice IC, House JI, Holland EA. 2005. Long-term sensitivity of soil carbon turnover to warming. Nature 433: 298–301.

Koricheva J, Gurevitch J, Mengersen K (Eds.). 2013. Handbook of Meta-analysis in Ecology and Evolution. Princeton University Press.

Kuznetsova A, Brockhoff PB, Christensen RHB. 2017. lmerTest Package: Tests in Linear Mixed Effects Models.

Lê S, Josse J, Husson F. 2008. FactoMineR : An *R* Package for Multivariate Analysis. Journal of Statistical Software 25.

Lenth R. 2019. emmeans: Estimated Marginal Means, aka Least-Squares Means.

Lind L, Harbicht A, Bergman E, Edwartz J, Eckstein RL. 2022. Effects of initial leaching for estimates of mass loss and microbial decomposition—Call for an increased nuance. Ecology and Evolution 12: e9118.

Liski J, Nissinen A, Erhard M, Taskinen O. 2003. Climatic effects on litter decomposition from arctic tundra to tropical rainforest. Global Change Biology 9: 575–584.

Luo C, Xu G, Chao Z, et al. 2010. Effect of warming and grazing on litter mass loss and temperature sensitivity of litter and dung mass loss on the Tibetan plateau. Global Change Biology 16: 1606–1617.

Marion GM, Henry GHR, Freckman DW, et al. 1997. Open-top designs for manipulating field temperature in high-latitude ecosystems. Global Change Biology 3: 20–32.

McCormack ML, Dickie IA, Eissenstat DM, et al. 2015. Redefining fine roots improves understanding of below-ground contributions to terrestrial biosphere processes. New Phytologist 207: 505–518.

Meentemeyer V. 1978. Macroclimate and Lignin Control of Litter Decomposition Rates. Ecology 59: 465–472.

Moise ERD, Henry HAL. 2014. Interactive responses of grass litter decomposition to warming, nitrogen addition and detritivore access in a temperate old field. OECOLOGIA 176: 1151–1160.

Mokany K, Raison RJ, Prokushkin AS. 2006. Critical analysis of root : shoot ratios in terrestrial biomes. Global Change Biology 12: 84–96.

Munir TM, Khadka B, Xu B, Strack M. 2017. Mineral nitrogen and phosphorus pools affected by water table lowering and warming in a boreal forested peatland. Ecohydrology 10: e1893.

Myers-Smith IH, Forbes BC, Wilmking M, et al. 2011. Shrub expansion in tundra ecosystems: dynamics, impacts and research priorities. Environmental Research Letters 6: 045509.

Nakagawa S, Lagisz M, O’Dea RE, et al. 2021. The orchard plot: Cultivating a forest plot for use in ecology, evolution, and beyond. Research Synthesis Methods 12: 4–12.

Parton WJ, Schimel DS, Cole CV, Ojima DS. 1987. Analysis of Factors Controlling Soil Organic Matter Levels in Great Plains Grasslands. Soil Science Society of America Journal 51: 1173–1179.

Pearson RG, Phillips SJ, Loranty MM, et al. 2013. Shifts in Arctic vegetation and associated feedbacks under climate change. Nature Climate Change 3: 673–677.

Poorter H, Niklas KJ, Reich PB, Oleksyn J, Poot P, Mommer L. 2012. Biomass allocation to leaves, stems and roots: meta-analyses of interspecific variation and environmental control. New Phytologist 193: 30–50.

Powers JS, Montgomery RA, Adair EC, et al. 2009. Decomposition in tropical forests: a pan-tropical study of the effects of litter type, litter placement and mesofaunal exclusion across a precipitation gradient. Journal of Ecology 97: 801–811.

Prescott CE. 2010. Litter decomposition: what controls it and how can we alter it to sequester more carbon in forest soils? Biogeochemistry 101: 133–149.

Rantanen M, Karpechko AY, Lipponen A, et al. 2022. The Arctic has warmed nearly four times faster than the globe since 1979. Communications Earth & Environment 3: 1–10.

Ren H, Qin J, Yan B, Alata, Baoyinhexige, Han G. 2018. Mass loss and nutrient dynamics during litter decomposition in response to warming and nitrogen addition in a desert steppe. FRONTIERS OF AGRICULTURAL SCIENCE AND ENGINEERING 5: 64–70.

Rey A, Pegoraro E, Jarvis PG. 2008. Carbon mineralization rates at different soil depths across a network of European forest sites (FORCAST). European Journal of Soil Science 59: 1049–1062.

Rohatgi A. 2021.Webplotdigitizer: Version 4.5.

Romero-Olivares A, Allison S, Treseder K. 2017. Decomposition of recalcitrant carbon under experimental warming in boreal forest. PLOS ONE 12.

Sarneel JM, Sundqvist MK, Molau U, Björkman MP, Alatalo JM. 2020. Decomposition rate and stabilization across six tundra vegetation types exposed to >20 years of warming. Science of The Total Environment 724: 138304.

Schimel JP. 2018. Life in Dry Soils: Effects of Drought on Soil Microbial Communities and Processes. Annual Review of Ecology, Evolution, and Systematics 49: 409–432.

Schimel DS, Braswell BH, Holland EA, et al. 1994. Climatic, edaphic, and biotic controls over storage and turnover of carbon in soils. Global Biogeochemical Cycles 8: 279–293.

Schuur E, Bockheim J, Canadell JG, et al. 2008. Vulnerability of Permafrost Carbon to Climate Change: Implications for the Global Carbon Cycle. BioScience 58: 701–714.

Seres A, Kröel-Dulay G, Szakálas J, et al. 2022. The response of litter decomposition to extreme drought modified by plant species, plant part, and soil depth in a temperate grassland. Ecology and Evolution 12: e9652.

Shaw M, Harte J. 2001. Control of litter decomposition in a subalpine meadow-sagebrush steppe ecotone under climate change. ECOLOGICAL APPLICATIONS 11: 1206–1223.

Shu M, Zhao Q, Li Z, Zhang L, Wang P, Hu S. 2019. Effects of global change factors and living roots on root litter decomposition in a Qinghai-Tibet alpine meadow. SCIENTIFIC REPORTS 9.

Sierra CA, Malghani S, Loescher HW. 2017. Interactions among temperature, moisture, and oxygen concentrations in controlling decomposition rates in a boreal forest soil. Biogeosciences 14: 703–710.

Sterne JAC, Egger M. 2001. Funnel plots for detecting bias in meta-analysis. Journal of Clinical Epidemiology 54: 1046–1055.

Sterne JAC, Gavaghan D, Egger M. 2000. Publication and related bias in meta-analysis: Power of statistical tests and prevalence in the literature. Journal of Clinical Epidemiology 53: 1119–1129.

Suseela V, Tharayil N, Xing B, Dukes JS. 2013. Labile compounds in plant litter reduce the sensitivity of decomposition to warming and altered precipitation. New Phytologist 200: 122–133.

Swift MJ, Heal OW, Anderson Jonathan Michael, Anderson J. M. 1979. Decomposition in Terrestrial Ecosystems. University of California Press.

Tarnocai C, Canadell JG, Schuur E a. G, Kuhry P, Mazhitova G, Zimov S. 2009. Soil organic carbon pools in the northern circumpolar permafrost region. Global Biogeochemical Cycles 23.

Thakur MP, Reich PB, Hobbie SE, et al. 2018. Reduced feeding activity of soil detritivores under warmer and drier conditions. Nature Climate Change 8: 75–78.

Thomas H, Myers-Smith I, Høye T, et al. 2023. Litter quality outweighs climate as a driver of decomposition across the tundra biome. Life Sciences.

Tingley MP, Huybers P. 2013. Recent temperature extremes at high northern latitudes unprecedented in the past 600 years. Nature 496: 201–205.

Viechtbauer W. 2010. Conducting Meta-Analyses in *R* with the metafor Package. Journal of Statistical Software 36.

Vogel A, Eisenhauer N, Weigelt A, Scherer-Lorenzen M. 2013. Plant diversity does not buffer drought effects on early-stage litter mass loss rates and microbial properties. Global Change Biology 19: 2795–2803.

Walter J, Hein R, Beierkuhnlein C, et al. 2013. Combined effects of multifactor climate change and land-use on decomposition in temperate grassland. SOIL BIOLOGY & BIOCHEMISTRY 60: 10–18.

Wang E, Cresswell H, Xu J, Jiang Q. 2009. Capacity of soils to buffer impact of climate variability and value of seasonal forecasts. Agricultural and Forest Meteorology 149: 38–50.

Wang P, Heijmans MM, Mommer L, van Ruijven J, Maximov TC, Berendse F. 2016. Belowground plant biomass allocation in tundra ecosystems and its relationship with temperature. Environmental Research Letters 11: 055003.

Wickham H, Chang W, Wickham MH. 2016. Package ‘ggplot2.’ Create Elegant Data Visualisations Using the Grammar of Graphics. Version 2: 1–189.

Wu Z, Dijkstra P, Koch GW, Peñuelas J, Hungate BA. 2011. Responses of terrestrial ecosystems to temperature and precipitation change: a meta-analysis of experimental manipulation. Global Change Biology 17: 927–942.

Xia M, Talhelm AF, Pregitzer KS. 2015. Fine roots are the dominant source of recalcitrant plant litter in sugar maple-dominated northern hardwood forests. New Phytologist 208: 715–726.

Yi C, Wei S, Hendrey G. 2014. Warming climate extends dryness-controlled areas of terrestrial carbon sequestration. Scientific Reports 4: 5472.

Zaller JG, Caldwell MM, Flint SD, Ballare CL, Scopel AL, Sala OE. 2009. Solar UVB and warming affect decomposition and earthworms in a fen ecosystem in Tierra del Fuego, Argentina. GLOBAL CHANGE BIOLOGY 15: 2493–2502.

Zhang D, Hui D, Luo Y, Zhou G. 2008. Rates of litter decomposition in terrestrial ecosystems: global patterns and controlling factors. Journal of Plant Ecology 1: 85–93.

## Websites

European Centre for Medium-Range Weather Forecasts (2022). European State of the Climate. Available at: https://climate.copernicus.eu/esotc/2022. Last accessed 29 January 2024.

National Centers for Environmental Information (2023). Global Drought Information System. Available at: https://www.ncei.noaa.gov/access/monitoring/monthly-report/global-drought/202309. Last accessed 29 January 2024

