## Supplemental Material for "Macro-environment strongly interacts with warming in a global analysis of decomposition"

### Supporting Information


**List of Figures** 2

**List of Tables** 3

**Literature screening process** 4

**Peer-reviewed literature included in the meta-analysis** 5

**Locations of open-top chamber warming experiments measuring standardised plant litter (tea) decomposition** 10

**Macro-environmental factors** 12

**The effect of experimental-induced warming on decomposition** 21

**Plant functional types and plant organ types interacting with the position of incubation (on soil surface, buried in the soil)** 22

### Supporting Information

#### List of Figures

***Figure S1*** *The literature screening process visualized as a preferred reporting items for systematic reviews and meta-analyses (PRISMA) flow diagram describing the number of screened studies (n) and exclusion rules in this meta-analysis.* ........ *4*

***Figure S2*** *(A) Global distribution of study sites coloured according to the four main macro-environmental classes derived from the principal component analysis. (B) Study sites plotted in a Whittaker Biome Diagram with dots for study sites coloured according to the four main macro-environmental classes.* ........ *14*

***Figure S3*** *Effects of experimental warming on plant litter decomposition. The pooled average decomposition standardised mean difference (SMD, Hedges' g; outlined circles) and 95% confidence intervals (black error bars) resulting from warming for the macro-environmental classes cold and dry (outlined circles), cold and wet (outlined squares), warm and dry (outlined diamonds), and warm and wet (outlined triangles) for the natural litter (blue, number of effect sizes k=527) and the standardised plant litter, separated into rooibos (red, k=57) and green tea (green, k=57). Each coloured dot is an individual effect size (non-outlined circles) with dot size representing its precision (the inverse of the standard error, larger points having greater influence on the model). Asterisks indicate that the overall pooled average SMD is significantly different from zero (**$p < 0.01$).* ........ *20*

***Figure S4*** *Impacts of experimentally induced changes in micro-environment on decomposition and its interaction with macro-environment. Effect of (A) degree of warming (i.e., absolute temperature difference between warmed and control plots, k=315); and (B) warming-induced changes in soil moisture with warming (i.e., difference between warmed and control plots in soil moisture, k=315) on decomposition SMD. Each coloured dot is an individual effect size (non-outlined circles) with dot size representing its precision (the inverse of the standard error, larger points having greater influence on the model). Asterisks indicate that the overall pooled average SMD is significantly different from zero. Solid lines indicate regression lines with shaded areas representing the 95%CI (***$p < 0.001$). Dashed lines indicate no significant relationship (n.s. = not significant).* ........ *21*

***Figure S5*** *Impact of warming methods on decomposition SMD. The pooled average decomposition standardised mean difference (SMD, Hedges' g; outlined circles) and 95% confidence intervals (black error bars) resulting from warming for the different experimental warming methods (see Table S1). Each coloured dot is an individual effect size (non-outlined circles) with dot size representing its precision (the inverse of the standard error, larger points having greater influence on the model). Letters indicate significant differences between the pooled average SMD of warming methods. Asterisks indicate a significant deviation of decomposition SMD from zero (*$p \leq 0.05$).* ........ *21*

***Figure S6*** *Differences in C:N ratio and warming effect on decomposition across plant functional types. (A) Plant functional types ranked based on carbon to nitrogen ratios (C:N ratios). Large, coloured points represent mean C:N ratios and small transparent dots individual plant species. (B) The pooled average decomposition standardized mean difference (SMD, Hedges' g, black outlined circles) and 95% confidence intervals (95%CI, black error bars) per plant functional type of natural litter and standardised plant litter combining data from above and below ground incubations. Different letters indicate differences in (A) mean C:N ratio and (B) decomposition SMD between the different plant functional litter types, as well as the standard material green and rooibos tea. Asterisks indicate that the overall pooled average SMD is significantly different from zero (*$p < 0.05$, **$p < 0.01$, ***$p < 0.001$).* ........ *22*

***Figure S7*** *Differences in ambient decomposability, measured as ambient mass loss rate per day (% $d^{-1}$), for the plant functional types and plant organs of natural plant litter and the standardised tea material (i.e., rooibos and green tea) for each of the four macro-environmental classes. Colours indicate the four macro-environmental classes of temperature (temp), precipitation (prec) and soil organic carbon (SOC) that are either high (▲) or low (▼), consistent with Figure 3 in the main text. Different letters indicate significant differences in decomposition SMD between plant functional types.* ........ *23*

### Supporting Information

#### List of Tables

***Table S1*** *Scientific research articles included in the meta-analysis, sorted by first author. The country of the study and used warming method (detailed information on the methods can be found in the original articles) of the reported/included study. Number of effect sizes per study (k), and sum of observations per study for the paired warming treatment and control.* ........ 5

***Table S2*** *Study sites in which standardised litter decomposition was measured in open-top chamber experiments. Observations per study are treatment replications in space and resulted in one effect size per site.* ........ 10

***Table S3*** *Correlation off the map-based macro-environmental climatic factors to the Principal component axes (PC1, PC2) together with the units and sources, including WorldClim2 = database of high spatial resolution global weather and climate data, SoilGrids = system for global digital soil mapping, CGIAR=Consortium of International Agricultural Research Centers, EarthEnv = Global, remote-sensing supported environmental layers for assessing status and trends in biodiversity, ecosystems, and climate, MODIS=Moderate Resolution Imaging Spectroradiometer.* ........ 12

***Table S4*** *Means and standard error (SE) of the map-based macro-environmental factors per macro-environmental class that are defined by the scores on the PCA axis and the correlation of these axis to climatic variables of temperature (temp), precipitation (prec),Table S4 Means and standard error (SE) of the map-based macro-environmental factors per macro-environmental class that are defined by the scores on the PCA axis and the correlation of these axis to climatic variables of temperature (temp), precipitation (prec), and soil organic carbon (SOC) that are either high (upward arrow) or low (downward arrow).* ........ 15

***Table S5*** *Results of single effects multivariate linear mixed-effects models for reported and measured macro-environmental factors with the standardised mean difference of decomposition (SMD) as dependent and measured or reported site-specific environmental factors as predictor. Values in bold indicate significant effect of the predictor on decomposition SMD (p ≤ 0.05). The number of effect sizes (k) used in the models, lower and upper bounds of the 95% confidence intervals, and heterogeneity explained by the model structure ($Q_M$) are reported.* ........ 18

***Table S6*** *Map-based macro-environmental results of single multivariate linear mixed-effects models with the standardised mean difference of decomposition (SMD) as dependent variable and the map-derived macro-environmental factors as predictor. Values in bold indicate significant effect of the predictor on decomposition SMD (p ≤ 0.05). The number of effect sizes (k) used in the models, lower and upper bounds of the 95% confidence intervals, and heterogeneity explained by the model structure ($Q_M$) are reported.* ........ 18

***Table S7*** *The impact of the four macro-environmental classes four macro-environmental classes distinguished by different combinations of high (▲) or low (▼) of temperature (temp), precipitation (prec) and soil organic carbon (SOC) and the natural and the standardised plant litter (i.e., green and rooibos tea) on the effect of warming on decomposition (SMD). Bold values indicate a significant effect of the macro-environmental class and litter type on SMD (p ≤ 0.05 or CI ≠ 0). Number of effect sizes (k), p-values, and 95%-confidence interval are shown.* ........ 20

***Table S8*** *The pooled average decomposition standardised mean difference (SMD) of different plant functional types of the natural litter and natural and the standardised plant litter (i.e., green and rooibos tea) with respect to the position of incubation (i.e., on soil surface, buried in the soil) as well as the number of effect sizes (k) for each category, the p-value and 95%-confidence interval describing whether the pooled average SMD significantly differs from zero (in bold, p ≤ 0.05). For forbs and nonvascular plants no reports of buried or root litter were available.* ........ 23

### Supporting Information

#### Literature screening process

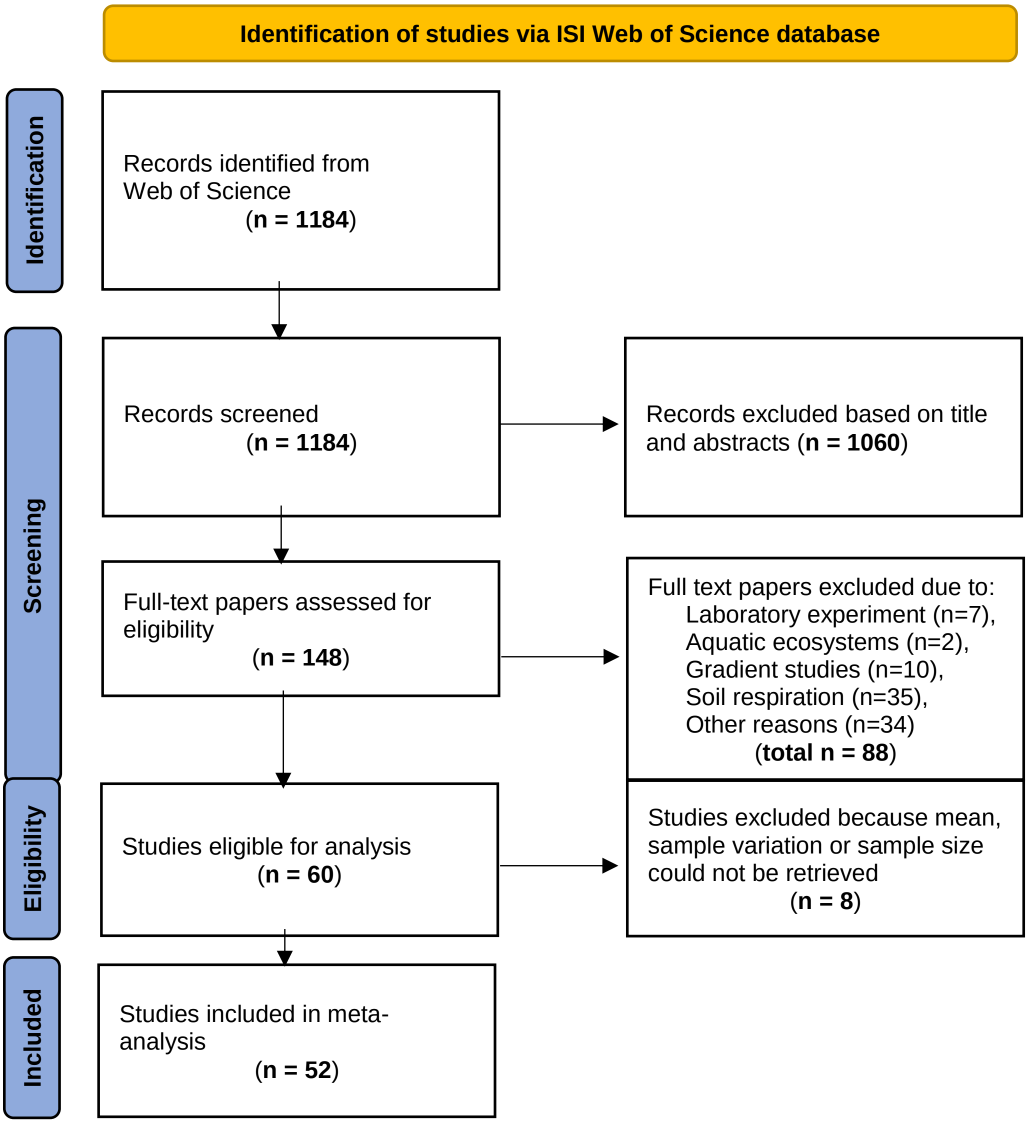


**Figure S1** The literature screening process visualized as a preferred reporting items for systematic reviews and meta-analyses (PRISMA) flow diagram describing the number of screened studies (n) and exclusion rules in this meta-analysis.

### Supporting Information

#### Peer-reviewed literature included in the meta-analysis

**Table S1** Scientific research articles included in the meta-analysis, sorted by first author. The country of the study and used warming method (detailed information on the methods can be found in the original articles) of the reported study. Number of effect sizes per study (k), and sum of observations from ambient vs warmed treatments per study for the paired warming treatment and control.

| Nr | Study | DOI | Country | Warming method | Effect sizes (*k*) | Obser-vations |
|---|---|---|---|---|---|---|
| 1 | Aerts et al., 2012 | 10.1007/s00442-012-2330-z | USA | Open-top chamber | 6 | 30 |
| 2 | Bélanger et al., 2023 | 10.3390/soilsystems7010014 | Kazakhstan | Heating cable | 24 | 480 |
| 3 | Berbeco et al., 2012 | 10.1007/s11104-012-1130-x | USA | Heating cable | 16 | 144 |
| 4 | Berdugo et al., 2021 | 10.1007/s10021-020-00599-0 | Spain | Open-top chamber | 12 | 57 |
| 5 | Bhuiyan et al., 2023 | 10.1016/j.scitotenv.2022.159683 | Finland | Open-top chamber | 50 | 300 |
| 6 | Blok et al., 2016 | 10.1007/s10021-015-9924-3 | Greenland | Open-top chamber | 6 | 36 |
| 7 | Blok et al., 2018 | 10.1111/gcb.14017 | Greenland | Open-top chamber | 4 | 20 |
| 8 | Bokhorst et al., 2010 | 10.1016/j.soilbio.2009.12.011 | Sweden | Infrared heater | 12 | 72 |
| 9 | Brigham et al., 2018 | 10.1080/15230430.2018.1494941 | Belgium | Open-top chamber | 4 | 16 |
| 10 | Carbognani et al., 2014 | 10.1007/s11104-013-1982-8 | Italy | Open-top chamber | 4 | 20 |
| 11 | Chen et al., 2008 | 10.2134/jeq2007.0266 | USA | Sunlit controlled-environment chamber | 2 | 78 |
| 12 | Cheng et al., 2010 | 10.1016/j.agee.2010.04.019 | Sweden | Infrared heater | 14 | 70 |
| 13 | Christiansen et al., 2017 | 10.1111/gcb.13362 | Greenland | Open-top chamber | 2 | 24 |
| 14 | Chuckran et al., 2020 | 10.1016/j.soilbio.2020.107799 | USA | Infrared heater | 6 | 120 |
| 15 | Cui et al., 2021 | 10.1007/s13157-021-01445-2 | China | Open-top chamber | 20 | 60 |
| 16 | De Long et al., 2016 | 10.1016/j.soilbio.2016.04.009 | Sweden | Open-top chamber | 6 | 60 |
| 17 | Gewirtzman et al., 2019 | 10.3389/fpls.2019.01097 | USA | Heating cable | 23 | 77 |
| 18 | Gong et al., 2015 | 10.1371/journal.pone.0116013 | China | Infrared heater | 6 | 36 |
| 19 | Han et al., 2019 | 10.3906/tar-1807-162 | South Korea | Infrared heater | 3 | 9 |
| 20 | Henry et al., 2015 | 10.1007/s11104-014-2346-8 | Canada | Infrared heater | 8 | 80 |
| 21 | Hong et al., 2021 | 10.1016/j.scitotenv.2020.142306 | China | Open-top chamber | 20 | 60 |
| 22 | Kasurinen et al., 2017 | 10.1007/s11104-016-3122-8 | Finland | Infrared heater | 2 | 28 |
| 23 | Li et al., 2022a | 10.1016/j.soilbio.2022.108716 | China | Heating cable | 14 | 140 |

| | | | | | | |
|---|---|---|---|---|---|---|
| 24 | Li et al., 2022b | 10.1093/jpe/rtac009 | China | Infrared heater | 1 | 8 |
| 25 | Liu et al., 2021 | 10.1007/s11104-020-04551-y | China | Infrared heater | 6 | 30 |
| 26 | Liu et al., 2022 | 10.1016/j.geoderma.2022.116139 | China | Heating cable | 8 | 160 |
| 27 | Lukas et al., 2018 | 10.1016/j.apsoil.2017.10.018 | Germany | Heating cable | 4 | 16 |
| 28 | Luo et al., 2010 | 10.1111/j.1365-2486.2009.02026.x | China | Infrared heater | 2 | 8 |
| 29 | Luo et al., 2023 | 10.1098/rspb.2023.0613 | China | Open-top chamber | 8 | 160 |
| 30 | McHale et al., 1998 | 10.1139/cjfr-28-9-1365 | USA | Heating cable | 12 | 108 |
| 31 | Moise et al., 2014 | 10.1007/s00442-014-3068-6 | Canada | Infrared heater | 4 | 40 |
| 32 | Morrison et al., 2019 | 10.1016/j.soilbio.2019.02.005 | USA | Heating cable | 1 | 10 |
| 33 | Prieto et al., 2019 | 10.1111/1365-2745.13168 | Spain | Open-top chamber | 2 | 20 |
| 34 | Remy et al., 2018 | 10.1007/s10021-017-0182-4 | Netherlands | Open-top chamber | 24 | 48 |
| 35 | Ren et al., 2018 | 10.15302/J-FASE-2017194 | China | Infrared heater | 5 | 30 |
| 36 | Robinson et al., 1995 | 10.2307/3545996 | Sweden; Norway:Svalbard | Open-topped polythene tents | 5 | 30 |
| 37 | Robinson et al., 1997 | 10.1046/j.1365-2486.1997.d01-133.x | Sweden; Norway:Svalbard | Open-topped polythene tents | 11 | 72 |
| 38 | Romero-Olivares et al., 2017 | 10.1371/journal.pone.0179674 | USA:Alaska | Open-top chamber | 4 | 20 |
| 39 | Rustad et al., 1998 | 10.2136/sssaj1998.03615995006200040031x | USA | Heating cable | 6 | 40 |
| 40 | Shaw et al., 2001 | 10.2307/3061022 | USA | Infrared heater | 34 | 176 |
| 41 | Shu et al., 2019 | 10.1038/s41598-019-53450-5 | China | Open-top chamber | 2 | 8 |
| 42 | Sjögersten et al., 2004 | 10.1023/B:PLSO.0000037044.63113.fe | Norway; Greenland | Open-top chamber | 60 | 300 |
| 43 | Sjögersten et al., 2012 | 10.1007/s10021-011-9514-y | Norway:Svalbard | Open-top chamber | 4 | 20 |
| 44 | Suseela et al., 2014 | 10.1016/j.soilbio.2014.03.022 | USA | Infrared heater | 6 | 36 |
| 45 | Walter et al., 2013 | 10.1016/j.soilbio.2013.01.018 | Germany | Infrared heater | 1 | 15 |
| 46 | Ward et al., 2015 | 10.1890/14-0292.1 | UK | Open-top chamber | 6 | 24 |
| 47 | Xu et al., 2012 | 10.1111/j.1365-2389.2012.01449.x | China | Heating cable | 9 | 36 |
| 48 | Ye et al., 2022 | 10.1016\j.soilbio.2022.108588 | China | Open-top chamber | 12 | 576 |
| 49 | Yin et al., 2022 | 10.1007/s00374-022-01639-8 | China | Heating cable | 4 | 32 |
| 50 | Yoshitake et al., 2021 | 10.1111/grs.12319 | Japan:Honshu | Infrared heater | 8 | 48 |
| 51 | Zaller et al., 2009 | 10.1111/j.1365-2486.2009.01970.x | Argentina:Tierra del Fuego | UVB filter film | 8 | 40 |
| 52 | Zhou et al., 2022 | 10.1093/jpe/rtac027 | China | Infrared heater | 2 | 16 |

1. Aerts R, Callaghan TV, Dorrepaal E, Van Logtestijn RSP, Cornelissen JHC. 2012. Seasonal climate manipulations have only minor effects on litter decomposition rates and N dynamics but strong effects on litter P dynamics of sub-arctic bog species. *Oecologia* 170: 809–819.
2. Bélanger N, Chaput-Richard C. 2023. Experimental warming of typically acidic and nutrient-poor boreal soils does not affect leaf-litter decomposition of temperate deciduous tree species. *SOIL SYSTEMS* 7.
3. Berbeco MR, Melillo JM, Orians CM. 2012. Soil warming accelerates decomposition of fine woody debris. *Plant and Soil* 356: 405–417.
4. Berdugo M, Mendoza-Aguilar DO, Rey A, *et al.* 2021. Litter Decomposition Rates of Biocrust-Forming Lichens Are Similar to Those of Vascular Plants and Are Affected by Warming. *Ecosystems* 24: 1531–1544.
5. Bhuiyan R, Makiranta P, Strakova P, *et al.* 2023. Fine-root biomass production and its contribution to organic matter accumulation in sedge fens under changing climate. *SCIENCE OF THE TOTAL ENVIRONMENT* 858.
6. Blok D, Elberling B, Michelsen A. 2016. Initial Stages of Tundra Shrub Litter Decomposition May Be Accelerated by Deeper Winter Snow But Slowed Down by Spring Warming. *Ecosystems* 19: 155–169.
7. Blok D, Faucherre S, Banyasz I, Rinnan R, Michelsen A, Elberling B. 2018. Contrasting above- and belowground organic matter decomposition and carbon and nitrogen dynamics in response to warming in High Arctic tundra. *Global Change Biology* 24: 2660–2672.
8. Bokhorst S, Bjerke JW, Melillo J, Callaghan TV, Phoenix GK. 2010. Impacts of extreme winter warming events on litter decomposition in a sub-Arctic heathland. *Soil Biology and Biochemistry* 42: 611–617.
9. Brigham LM, Esch EH, Kopp CW, Cleland EE. 2018. Warming and shrub encroachment decrease decomposition in arid alpine and subalpine ecosystems. *Arctic, Antarctic, and Alpine Research* 50: e1494941.
10. Carbognani M, Petraglia A, Tomaselli M. 2014. Warming effects and plant trait control on the early-decomposition in alpine snowbeds. *Plant and Soil* 376: 277–290.
11. Chen H, Rygiewicz PT, Johnson MG, Harmon ME, Tian H, Tang JW. 2008. Chemistry and Long-Term Decomposition of Roots of Douglas-Fir Grown under Elevated Atmospheric Carbon Dioxide and Warming Conditions. *Journal of Environmental Quality* 37: 1327–1336.
12. Cheng X, Luo Y, Su B, *et al.* 2010. Experimental warming and clipping altered litter carbon and nitrogen dynamics in a tallgrass prairie. *Agriculture, Ecosystems & Environment* 138: 206–213.
13. Christiansen CT, Haugwitz MS, Priemé A, *et al.* 2017. Enhanced summer warming reduces fungal decomposer diversity and litter mass loss more strongly in dry than in wet tundra. *Global Change Biology* 23: 406–420.
14. Chuckran PF, Reibold R, Throop HL, Reed SC. 2020. Multiple mechanisms determine the effect of warming on plant litter decomposition in a dryland. *Soil Biology and Biochemistry* 145: 107799.
15. Cui W, Mao Y, Tian K, Wang H. 2021. A Comparative Study of Manipulative and Natural Temperature Increases in Controlling Wetland Plant Litter Decomposition. *Wetlands* 41: 48.
16. De Long JR, Dorrepaal E, Kardol P, Nilsson M-C, Teuber LM, Wardle DA. 2016. Understory plant functional groups and litter species identity are stronger drivers of litter decomposition than warming along a boreal forest post-fire successional gradient. *Soil Biology and Biochemistry* 98: 159–170.
17. Gewirtzman J, Tang J, Melillo JM, *et al.* 2019. Soil Warming Accelerates Biogeochemical Silica Cycling in a Temperate Forest. *Frontiers in Plant Science* 10: 1097.
18. Gong S, Guo R, Zhang T, Guo J. 2015. Warming and Nitrogen Addition Increase Litter Decomposition in a Temperate Meadow Ecosystem (M Schädler, Ed.). *PLOS ONE* 10: e0116013.
19. Han SH, Kim S, Chang H, Li G. 2019. Increased soil temperature stimulates changes in carbon, nitrogen, and mass loss in the fine roots of Pinus koraiensis under experimental warming and drought. *TURKISH JOURNAL OF AGRICULTURE AND FORESTRY* 43: 80–87.
20. Henry HAL, Moise ERD. 2015. Grass litter responses to warming and N addition: temporal variation in the contributions of litter quality and environmental effects to decomposition. *Plant and Soil* 389: 35–43.

21. Hong J, Lu X, Ma X, Wang X. 2021. Five-year study on the effects of warming and plant litter quality on litter decomposition rate in a Tibetan alpine grassland. *Science of The Total Environment* 750: 142306.
22. Kasurinen A, Silfver T, Rousi M, Mikola J. 2017. Warming and ozone exposure effects on silver birch (Betula pendula Roth) leaf litter quality, microbial growth and decomposition. *Plant and Soil* 414: 127–142.
23. Li A, Fan Y, Chen S, Song H, Lin C, Yang Y. 2022. Soil warming did not enhance leaf litter decomposition in two subtropical forests. *SOIL BIOLOGY & BIOCHEMISTRY* 170.
24. Li B, Lv W, Sun J, *et al.* 2022. Warming and grazing enhance litter decomposition and nutrient release independent of litter quality in an alpine meadow. *JOURNAL OF PLANT ECOLOGY* 15: 977–990.
25. Liu X, Chen S, Li X, *et al.* 2022. Soil warming delays leaf litter decomposition but exerts no effect on litter nutrient release in a subtropical natural forest over 450 days. *GEODERMA* 427.
26. Liu H, Lin L, Wang H, *et al.* 2021. Simulating warmer and drier climate increases root production but decreases root decomposition in an alpine grassland on the Tibetan plateau. *Plant and Soil* 458: 59–73.
27. Lukas S, Abbas SJ, Kössler P, Karlovsky P, Potthoff M, Joergensen RG. 2018. Fungal plant pathogens on inoculated maize leaves in a simulated soil warming experiment. *Applied Soil Ecology* 124: 75–82.
28. Luo B, Huang M, Wang W, *et al.* 2023. Ant nests increase litter decomposition to mitigate the negative effect of warming in an alpine grassland ecosystem. *PROCEEDINGS OF THE ROYAL SOCIETY B-BIOLOGICAL SCIENCES* 290.
29. Luo C, Xu G, Chao Z, *et al.* 2010. Effect of warming and grazing on litter mass loss and temperature sensitivity of litter and dung mass loss on the Tibetan plateau. *Global Change Biology* 16: 1606–1617.
30. McHale PJ, Mitchell MJ, Bowles FP. 1998. Soil warming in a northern hardwood forest: trace gas fluxes and leaf litter decomposition. *Canadian Journal of Forest Research* 28: 1365–1372.
31. Moise ERD, Henry HAL. 2014. Interactive responses of grass litter decomposition to warming, nitrogen addition and detritivore access in a temperate old field. *Oecologia* 176: 1151–1160.
32. Morrison EW, Pringle A, Van Diepen LTA, Grandy AS, Melillo JM, Frey SD. 2019. Warming alters fungal communities and litter chemistry with implications for soil carbon stocks. *Soil Biology and Biochemistry* 132: 120–130.
33. Prieto I, Almagro M, Bastida F, Querejeta JI. 2019. Altered leaf litter quality exacerbates the negative impact of climate change on decomposition (P Kardol, Ed.). *Journal of Ecology* 107: 2364–2382.
34. Remy E, Wuyts K, Van Nevel L, De Smedt P, Boeckx P, Verheyen K. 2018. Driving Factors Behind Litter Decomposition and Nutrient Release at Temperate Forest Edges. *Ecosystems* 21: 755–771.
35. Ren H, Qin J, Yan B, Alata, Baoyinhexige, Han G. 2018. Mass loss and nutrient dynamics during litter decomposition in response to warming and nitrogen addition in a desert steppe. *Frontiers of Agricultural Science and Engineering* 5: 64.
36. Robinson CH, Michelsen A, Lee JA, *et al.* 1997. Elevated atmospheric $CO_2$ affects decomposition of *Festuca vivipara* (L.) Sm. litter and roots in experiments simulating environmental change in two contrasting arctic ecosystems. *Global Change Biology* 3: 37–49.
37. Robinson CH, Wookey PA, Parsons AN, *et al.* 1995. Responses of Plant Litter Decomposition and Nitrogen Mineralisation to Simulated Environmental Change in a High Arctic Polar Semi-Desert and a Subarctic Dwarf Shrub Heath. *Oikos* 74: 503.
38. Romero-Olivares AL, Allison SD, Treseder KK. 2017. Decomposition of recalcitrant carbon under experimental warming in boreal forest (D Hui, Ed.). *PLOS ONE* 12: e0179674.
39. Rustad LE, Fernandez IJ. 1998. Soil Warming: Consequences for Foliar Litter Decay in a Spruce-Fir Forest in Maine, USA. *Soil Science Society of America Journal* 62: 1072–1080.
40. Shaw MR, Harte J. 2001. Control of Litter Decomposition in a Subalpine Meadow-Sagebrush Steppe Ecotone under Climate Change. *Ecological Applications* 11: 1206.
41. Shu M, Zhao Q, Li Z, Zhang L, Wang P, Hu S. 2019. Effects of global change factors and living roots on root litter decomposition in a Qinghai-Tibet alpine meadow. *Scientific Reports* 9: 16924.

42. Sjögersten S, Van Der Wal R, Woodin SJ. 2012. Impacts of Grazing and Climate Warming on C Pools and Decomposition Rates in Arctic Environments. *Ecosystems* 15: 349–362.
43. Sjögersten S, Wookey PA. 2004. Decomposition of mountain birch leaf litter at the forest-tundra ecotone in the Fennoscandian mountains in relation to climate and soil conditions. *Plant and Soil* 262: 215–227.
44. Suseela V, Tharayil N, Xing B, Dukes JS. 2014. Warming alters potential enzyme activity but precipitation regulates chemical transformations in grass litter exposed to simulated climatic changes. *Soil Biology and Biochemistry* 75: 102–112.
45. Walter J, Hein R, Beierkuhnlein C, *et al.* 2013. Combined effects of multifactor climate change and land-use on decomposition in temperate grassland. *Soil Biology and Biochemistry* 60: 10–18.
46. Ward SE, Orwin KH, Ostle NJ, *et al.* 2015. Vegetation exerts a greater control on litter decomposition than climate warming in peatlands. *Ecology* 96: 113–123.
47. Xu ZF, Pu XZ, Yin HJ, Zhao CZ, Liu Q, Wu FZ. 2012. Warming effects on the early decomposition of three litter types, Eastern Tibetan Plateau, China. *European Journal of Soil Science* 63: 360–367.
48. Ye C, Wang Y, Yan X, Guo H. 2022. Predominant role of air warming in regulating litter decomposition in a Tibetan alpine meadow: A multi-factor global change experiment. *SOIL BIOLOGY & BIOCHEMISTRY* 167.
49. Yin R, Qin W, Zhao H, Wang X, Cao G, Zhu B. 2022. Climate warming in an alpine meadow: differential responses of soil faunal vs. microbial effects on litter decomposition. *BIOLOGY AND FERTILITY OF SOILS* 58: 509–514.
50. Yoshitake S, Suminokura N, Ohtsuka T, Koizumi H. 2021. Composite effects of temperature increase and snow cover change on litter decomposition and microbial community in cool-temperate grassland. *Grassland Science* 67: 315–327.
51. Zaller JG, Caldwell MM, Flint SD, Ballaré CL, Scopel AL, Sala OE. 2009. Solar UVB and warming affect decomposition and earthworms in a fen ecosystem in Tierra del Fuego, Argentina. *Global Change Biology* 15: 2493–2502.
52. Zhou Y, Lv W-W, Wang S-P, *et al.* 2022. Additive effects of warming and grazing on fine-root decomposition and loss of nutrients in an alpine meadow. *JOURNAL OF PLANT ECOLOGY* 15: 1273–1284.

### Supporting Information

#### Locations of open-top chamber warming experiments measuring standardised plant litter (tea) decomposition

**Table S2** Study sites in which standardised litter decomposition was measured in open-top chamber experiments. Observations per study are treatment replications in space and resulted in one effect size per site.

| Nr | Site_ID | Site name | Country | Observations |
|---|---|---|---|---|
| 1 | ATA_1 | Anchorage Island | Greenland | 5 |
| 2 | AUS_1 | Australia | Australia | 4 |
| 3 | CAN_1 | Common garden | Canada | 12 |
| 4 | CAN_2 | Drained peatland | Canada | 6 |
| 5 | CAN_3 | Kluane Elevation Transect 1 | Canada | 4 |
| 6 | CAN_4 | Kluane Elevation Transect 10 | Canada | 4 |
| 7 | CAN_5 | Kluane Elevation Transect 4 | Canada | 4 |
| 8 | CAN_6 | Kluane Elevation Transect 7 | Canada | 3 |
| 9 | CAN_7 | Plot B_dry | Canada | 4 |
| 10 | CAN_9 | Pristine peatland | Canada | 6 |
| 11 | CHN_1 | China meadow | China | 18 |
| 12 | CHN_2 | China mountain | China | 19 |
| 13 | CHN_3 | China swamp | China | 18 |
| 14 | CHN_4 | National Field Observation and Research Station of Agro-ecosystems | China | 9 |
| 15 | ESP_1 | Santa Olla | Spain | 6 |
| 16 | GRL_1 | High_altitude - mesic mixed shrub tundra | Greenland | 6 |
| 17 | GRL_2 | Low_altitude - mesic mixed shrub tundra | Greenland | 6 |
| 18 | ISL_1 | Audkuluheidi | Iceland | 20 |
| 19 | ISL_2 | Thingvellir | Iceland | 19 |
| 20 | ITA_1 | Moss-snowbed | Italy | 5 |
| 21 | ITA_2 | Shrub-snowbed | Italy | 5 |
| 22 | ITA_3 | Po Valley | Italy | 5 |
| 23 | ITA_4 | Northern Apennine | Italy | 5 |
| 24 | JPN_1 | NKM2601 | Japan:Honshu | 10 |
| 25 | JPN_2 | Sapporo | Japan:Hokkaido | 8 |
| 26 | JPN_3 | SGDG | Japan:Honshu | 8 |
| 27 | NOR_1 | ITEX site Finse | Norway | 17 |
| 28 | NOR_2 | Gudmedalen - low elevation | Norway | 7 |
| 29 | NOR_3 | Kongsvoll Lower dry tundra | Norway | 5 |
| 30 | NOR_4 | Kongsvoll Lower mesic tundra | Norway | 4 |
| 31 | NOR_5 | Kongsvoll Upper mesic tundra | Norway | 5 |
| 32 | RUS_1 | OTC experimental site, Eriophorum-Sphagnum bog | Russia | 8 |
| 33 | RUS_2 | OTC experimental site, Sphagnum bog | Russia | 8 |
| 34 | SAU_1 | Saudi Arabia | Saudi Arabia | 10 |
| 35 | SJM_1 | Endalen - Cassiope heath | Norway:Svalbard | 19 |
| 36 | SJM_2 | Endalen - Dryas heath | Norway:Svalbard | 18 |
| 37 | SJM_3 | Endalen - Moss tundra | Norway:Svalbard | 19 |
| 38 | SJM_4 | Endalen - Snowbed community | Norway:Svalbard | 10 |
| 39 | SJM_5 | Svalbard_mesic | Norway:Svalbard | 12 |

| | | | | |
|---|---|---|---|---|
| 40 | SJM_6 | Svalbard_moist | Norway:Svalbard | 14 |
| 41 | SJM_7 | Svalbard_wet | Norway:Svalbard | 14 |
| 42 | SWE_1 | Abisko | Sweden | 5 |
| 43 | SWE_2 | Latnajaure – Mesic meadow | Sweden | 9 |
| 44 | SWE_3 | Latnajaure – Dry heath | Sweden | 3 |
| 45 | SWE_4 | Latnajaure – Dry meadow | Sweden | 5 |
| 46 | SWE_5 | Latnajaure – Wet meadow | Sweden | 4 |
| 47 | SWE_6 | Latnajaure – Tussock tundra | Sweden | 5 |
| 48 | SWE_7 | Latnajaure – Wet meadow | Sweden | 5 |
| 49 | USA_1 | Atqasuk ITEX Dry Site | USA:Alaska | 6 |
| 50 | USA_2 | Atqasuk ITEX Wet Site | USA:Alaska | 6 |
| 51 | USA_3 | Barrow ITEX Dry Site | USA:Alaska | 6 |
| 52 | USA_4 | Barrow ITEX Wet Site | USA:Alaska | 6 |
| 53 | ZAF_1 | Cathedral Peak - grassland052rburn | South Africa | 4 |
| 54 | ZAF_2 | Cathedral Peak - grassland0annual | South Africa | 4 |
| 55 | ZAF_3 | Cathedral Peak - grassland0biennual | South Africa | 4 |
| 56 | ZAF_4 | Cathedral Peak - grassland0noburn | South Africa | 3 |
| 57 | ZAF_5 | Cathedral Peak - grassland0slope | South Africa | 4 |

#### Macro-environmental factors

**Table S3** Correlation off the map-based macro-environmental climatic factors to the Principal component axes (PC1, PC2) together with the units and sources, including WorldClim2 = database of high spatial resolution global weather and climate data, SoilGrids = system for global digital soil mapping, CGIAR=Consortium of International Agricultural Research Centers, EarthEnv = Global, remote-sensing supported environmental layers for assessing status and trends in biodiversity, ecosystems, and climate, MODIS=Moderate Resolution Imaging Spectroradiometer.

| **Variables** | **Correlation coefficients** | | **Unit** | **Global climate layer** | **Source** |
|---|---|---|---|---|---|
| | **PCA1** | **PCA2** | | | |
| **Temperature** | | | | | |
| Annual Mean Temperature | 0.89 | 0.25 | °C | WorldClim2 | |
| Max Temperature of Warmest Month | 0.86 | 0.09 | °C | WorldClim2 | |
| Air temperature isothermality | 0.64 | -0.19 | unitless | WorldClim2 | |
| Mean Diurnal Range | 0.56 | -0.35 | °C | WorldClim2 | |
| Mean Temperature of Coldest Quarter | 0.81 | 0.28 | °C | WorldClim2 | |
| Mean Temperature of Driest Quarter | 0.56 | 0.12 | °C | WorldClim2 | |
| Mean Temperature of Warmest Quarter | 0.81 | 0.21 | °C | WorldClim2 | |
| Mean Temperature of Wettest Quarter | 0.57 | 0.15 | °C | WorldClim2 | |
| Min Temperature of Coldest Month | 0.68 | 0.33 | °C | WorldClim2 | |
| Annual Temperature Range | 0.08 | -0.26 | °C | WorldClim2 | |
| Temperature Seasonality | -0.19 | -0.15 | °C | WorldClim2 | |
| Mean Temperature During Incubation Period | 0.61 | 0.27 | °C | WorldClim2 | |
| **Precipitation** | | | | | |
| Annual Precipitation | 0.46 | 0.77 | mm | WorldClim2 | |
| Precipitation of Coldest Quarter | 0.20 | 0.82 | mm | WorldClim2 | |
| Precipitation of Driest Month | 0.19 | 0.87 | mm | WorldClim2 | |
| Precipitation of Driest Quarter | 0.24 | 0.88 | mm | WorldClim2 | |
| Precipitation of Warmest Quarter | 0.40 | 0.39 | mm | WorldClim2 | |
| Precipitation of Wettest Month | 0.51 | 0.41 | mm | WorldClim2 | |

| | | | | | |
|---|---|---|---|---|---|
| Precipitation of Wettest Quarter | 0.51 | 0.46 | mm | WorldClim2 | |
| Precipitation Seasonality | 0.13 | -0.63 | unitless | WorldClim2 | |
| Sum Precipitation During Incubation Period | -0.01 | 0.32 | mm | WorldClim2 | |
| **Soil** | | | | | |
| Bulk density at 5 cm depth | 0.73 | -0.21 | cg cm-3 | SoilGrids | https://www.soilgrids.org |
| SOC Content at 5 cm depth | -0.78 | 0.29 | dg kg-1 | SoilGrids | https://www.soilgrids.org |
| SOC Density at 5 cm depth | -0.73 | 0.33 | dg kg-1 | SoilGrids | https://www.soilgrids.org |
| SOC Stock 0-5 cm depth | -0.49 | 0.57 | kg m² | SoilGrids | https://www.soilgrids.org |
| Sum of Total Nitrogen at 5 cm depth | -0.53 | 0.62 | cg kg-1 | SoilGrids 2.0 | https://www.soilgrids.org |
| Sum of Total Nitrogen at 15 cm depth | -0.76 | 0.21 | cg kg-1 | SoilGrids 2.0 | https://www.soilgrids.org |
| Sum of Total Nitrogen at 30 cm depth | -0.76 | 0.08 | cg kg-1 | SoilGrids 2.0 | https://www.soilgrids.org |
| **Other** | | | | | |
| Annal Mean Solar Radiation | 0.77 | -0.35 | kJ/(m² day) | WorldClim2 | |
| Aridity Index | -0.23 | 0.77 | AI Value | CGIAR | http://www.cgiar-csi.org/data/global-aridity-and-pet-database |
| Aspect Cosine | 0.06 | -0.15 | degree | TopoMed | https://www.earthenv.org/topography |
| Aspect Sine | -0.07 | 0.34 | degree | TopoMed | https://www.earthenv.org/topography |
| Cover Barren | -0.53 | -0.19 | % (0-100) | Concensus | https://www.earthenv.org/landcover |
| Cover Cultivated | 0.48 | -0.31 | % (0-100) | Concensus | https://www.earthenv.org/landcover |
| Cover Deciduous Broadleaf Trees | 0.09 | 0.56 | % (0-100) | Concensus | https://www.earthenv.org/landcover |
| Cover Evergreen Broadleaf Trees | 0.14 | 0.22 | % (0-100) | Concensus | https://www.earthenv.org/landcover |
| Cover Evergreen Needleleaf Trees | -0.02 | 0.16 | % (0-100) | Concensus | https://www.earthenv.org/landcover |
| Cover Herbaceous | 0.01 | -0.54 | % (0-100) | Concensus | https://www.earthenv.org/landcover |
| Cover Regularly Flooded | -0.17 | 0.03 | % (0-100) | Concensus | https://www.earthenv.org/landcover |
| Cover Shrubs | -0.20 | -0.06 | % (0-100) | Concensus | https://www.earthenv.org/landcover |
| Eastness | -0.09 | 0.11 | index (-1 to 1) | TopoMed | https://www.earthenv.org/topography |
| Elevation | 0.15 | -0.56 | meters | TopoMed | https://www.earthenv.org/topography |
| Fraction Photosynthetically Active Radiation (fPAR) | 0.54 | 0.65 | Fpar fraction | MODIS | https://explorer.earthengine.google.com/#detail/MODIS%2F006%2FMCD15A3H |
| Soil water capacity at 5 cm depth | -0.56 | 0.05 | % | SoilGrids | https://www.soilgrids.org |

| | | | | | |
|---|---|---|---|---|---|
| Northness | 0.28 | -0.14 | index (-1 to 1) | TopoMed | https://www.earthenv.org/topography |
| Potential Evapotranspiration (PET) | 0.88 | -0.29 | PET Value (mm) | CGIAR | http://www.cgiar-csi.org/data/global-aridity-and-pet-database |
| Saturated Water Content 5 cm depth | -0.74 | 0.16 | % | SoilGrids | https://www.soilgrids.org |
| Soil pH (water) at 5 cm depth | 0.34 | -0.78 | pH x 10 | SoilGrids | https://www.soilgrids.org |

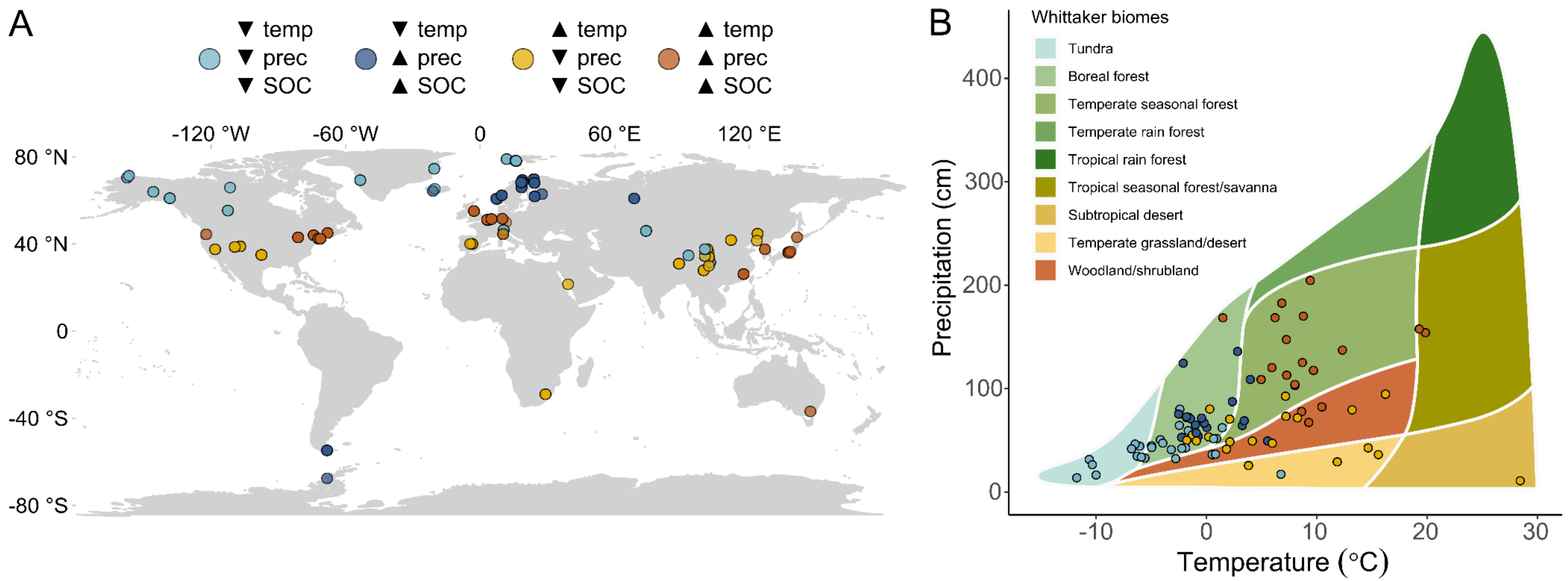


**Figure S2** (**A**) Global distribution of study sites coloured according to the four main macro-environmental classes derived from the principal component analysis. (**B**) Study sites plotted in a Whittaker Biome Diagram with dots for study sites coloured according to the four main macro-environmental classes.

**Table S4** Means and standard error (SE) of the map-based macro-environmental factors per macro-environmental class that are defined by the scores on the PCA axis and the correlation of these axis to climatic variables of temperature (temp), precipitation (prec),Table S4 Means and standard error (SE) of the map-based macro-environmental factors per macro-environmental class that are defined by the scores on the PCA axis and the correlation of these axis to climatic variables of temperature (temp), precipitation (prec), and soil organic carbon (SOC) that are either high (upward arrow) or low (downward arrow).

| **Variables** | **Unit** | **▲ temp ▲ prec ▼ SOC** | | **▲ temp ▼ prec ▼ SOC** | | **▼ temp ▲ prec ▲ SOC** | | **▼ temp ▼ prec ▲ SOC** | |
|---|---|---|---|---|---|---|---|---|---|
| | | mean | SE | mean | SE | mean | SE | mean | SE |
| **Temperature** | | | | | | | | | |
| Annual Mean Temperature | °C | 9.3 | ± 0.3 | 5.6 | ± 0.5 | 0.3 | ± 0.2 | -2.2 | ± 0.4 |
| Max Temperature of Warmest Month | °C | 24.6 | ± 0.3 | 22.8 | ± 0.5 | 15.5 | ± 0.2 | 14.7 | ± 0.5 |
| Isothermality | unitless | 31.1 | ± 0.3 | 37.8 | ± 0.4 | 25.4 | ± 0.3 | 23.7 | ± 0.5 |
| Mean Diurnal Range | °C | 9.8 | ± 0.1 | 12.9 | ± 0.1 | 7.4 | ± 0.1 | 8.0 | ± 0.2 |
| Mean Temperature of Coldest Quarter | °C | -0.9 | ± 0.5 | -4.4 | ± 0.6 | -9.1 | ± 0.3 | -13.5 | ± 0.4 |
| Mean Temperature of Driest Quarter | °C | 3.0 | ± 0.7 | 1.0 | ± 0.8 | -3.9 | ± 0.4 | -4.4 | ± 1.0 |
| Mean Temperature of Warmest Quarter | °C | 19.2 | ± 0.3 | 15.2 | ± 0.5 | 10.8 | ± 0.2 | 9.5 | ± 0.5 |
| Mean Temperature of Wettest Quarter | °C | 13.2 | ± 0.5 | 12.5 | ± 0.4 | 8.0 | ± 0.4 | 3.7 | ± 0.6 |
| Min Temperature of Coldest Month | °C | -7.5 | ± 0.6 | -12.2 | ± 0.6 | -14.2 | ± 0.3 | -19.1 | ± 0.4 |
| Annual Temperature Range | °C | 32.1 | ± 0.5 | 35.0 | ± 0.5 | 29.7 | ± 0.5 | 33.8 | ± 0.6 |
| Temperature Seasonality | °C | 819.1 | ± 15.0 | 799.5 | ± 17.5 | 807.1 | ± 15.0 | 945.5 | ± 19.8 |
| Mean Temperature during Incubation Period | °C | 11.1 | ± 0.5 | 6.7 | ± 0.6 | 1.4 | ± 0.4 | 2.4 | ± 0.5 |
| **Precipitation** | | | | | | | | | |
| Annual Precipitation | mm | 1172.4 | ± 24.4 | 554.2 | ± 15.3 | 642.1 | ± 13.7 | 357.5 | ± 13.1 |
| Precipitation of Coldest Quarter | mm | 241.6 | ± 7.1 | 67.1 | ± 4.2 | 141.6 | ± 5.5 | 58.5 | ± 2.7 |
| Precipitation of Driest Month | mm | 61.0 | ± 1.3 | 13.4 | ± 1.0 | 31.5 | ± 0.7 | 11.0 | ± 0.7 |
| Precipitation of Driest Quarter | mm | 204.4 | ± 3.7 | 49.9 | ± 3.2 | 103.4 | ± 2.2 | 42.6 | ± 2.2 |
| Precipitation of Warmest Quarter | mm | 337.4 | ± 12.4 | 224.4 | ± 8.6 | 208.9 | ± 2.6 | 140.9 | ± 7.2 |

| | | | | | | | | | | | | | |
|---|---|---|---|---|---|---|---|---|---|---|---|---|---|
| Precipitation of Wettest Month | mm | 142.5 | ± | 5.8 | 95.7 | ± | 2.9 | 85.1 | ± | 1.4 | 57.0 | ± | 2.6 |
| Precipitation of Wettest Quarter | mm | 399.8 | ± | 15.7 | 250.3 | ± | 7.4 | 228.1 | ± | 3.9 | 148.1 | ± | 6.9 |
| Precipitation Seasonality | unitless | 23.9 | ± | 1.7 | 63.5 | ± | 2.7 | 32.9 | ± | 0.6 | 47.4 | ± | 2.2 |
| Sum Precipitation during Incubation Period | mm | 820069.8 | ± | 56210.0 | 490908.5 | ± | 34505.2 | 912969.4 | ± | 47987.0 | 367516.6 | ± | 35516.5 |
| **Soil** | | | | | | | | | | | | | |
| Bulk density at 5 cm depth | cg $cm^{-3}$ | 905.0 | ± | 17.6 | 1070.4 | ± | 20.9 | 504.9 | ± | 12.4 | 736.7 | ± | 11.1 |
| SOC Content at 5 cm depth | dg $kg^{-1}$ | 78.9 | ± | 3.8 | 48.1 | ± | 2.1 | 142.8 | ± | 4.3 | 132.3 | ± | 3.8 |
| SOC Density at 5 cm depth | dg $kg^{-1}$ | 620.9 | ± | 19.3 | 447.4 | ± | 15.4 | 783.2 | ± | 8.8 | 748.9 | ± | 10.3 |
| SOC Stock 0-5 cm depth | kg $m^2$ | 41.2 | ± | 1.3 | 25.2 | ± | 0.8 | 38.1 | ± | 0.5 | 42.7 | ± | 0.7 |
| Sum of Total Nitrogen at 5 cm depth | cg $kg^{-1}$ | 8776.1 | ± | 321.6 | 4561.7 | ± | 175.2 | 9632.6 | ± | 103.5 | 7817.3 | ± | 200.2 |
| Sum of Total Nitrogen at 15 cm depth | cg $kg^{-1}$ | 3023.5 | ± | 78.2 | 2220.4 | ± | 61.1 | 5676.7 | ± | 163.6 | 5483.2 | ± | 191.8 |
| Sum of Total Nitrogen at 30 cm depth | cg $kg^{-1}$ | 2007.3 | ± | 44.4 | 1639.6 | ± | 39.6 | 3506.9 | ± | 112.3 | 4508.9 | ± | 165.8 |
| **Other** | | | | | | | | | | | | | |
| Annual Mean Solar Radiation | kJ/($m^2$ day) | 12532.0 | ± | 124.6 | 15999.4 | ± | 107.2 | 8170.1 | ± | 44.4 | 10200.2 | ± | 272.5 |
| Aridity Index | AI Value | 12066.2 | ± | 305.5 | 4484.7 | ± | 137.5 | 12164.9 | ± | 285.0 | 6978.2 | ± | 310.7 |
| Aspect Cosine | degree | 0.1 | ± | 0.0 | 0.0 | ± | 0.1 | 0.0 | ± | 0.1 | 0.3 | ± | 0.1 |
| Aspect Sine | degree | 0.2 | ± | 0.0 | -0.2 | ± | 0.0 | -0.1 | ± | 0.0 | -0.1 | ± | 0.0 |
| Cover Barren | % (0-100) | 1.8 | ± | 0.4 | 5.3 | ± | 1.0 | 12.7 | ± | 1.3 | 26.2 | ± | 1.9 |
| Cover Cultivated | % (0-100) | 12.1 | ± | 1.5 | 26.1 | ± | 2.1 | 0.2 | ± | 0.1 | 3.7 | ± | 0.9 |
| Cover Deciduous Broadleaf Trees | % (0-100) | 23.8 | ± | 2.1 | 1.5 | ± | 0.2 | 5.9 | ± | 0.9 | 1.4 | ± | 0.3 |
| Cover Evergreen Broadleaf Trees | % (0-100) | 2.3 | ± | 0.5 | 0.0 | ± | 0.0 | 1.9 | ± | 0.7 | 0.0 | ± | 0.0 |
| Cover Evergreen Needleleaf Trees | % (0-100) | 6.5 | ± | 1.0 | 12.0 | ± | 1.8 | 17.2 | ± | 2.0 | 2.0 | ± | 0.3 |
| Cover Herbaceous | % (0-100) | 4.0 | ± | 1.4 | 40.8 | ± | 2.0 | 8.3 | ± | 0.9 | 24.1 | ± | 2.3 |
| Cover Regularly Flooded | % (0-100) | 0.0 | ± | 0.0 | 0.0 | ± | 0.0 | 4.1 | ± | 1.3 | 0.5 | ± | 0.2 |
| Cover Shrubs | % (0-100) | 0.1 | ± | 0.1 | 6.1 | ± | 1.1 | 19.7 | ± | 1.1 | 5.2 | ± | 1.2 |
| Eastness | index (-1 to 1) | 0.0 | ± | 0.0 | 0.0 | ± | 0.0 | 0.1 | ± | 0.0 | 0.0 | ± | 0.0 |
| Elevation | meters | 348.8 | ± | 31.2 | 2585.0 | ± | 111.3 | 436.8 | ± | 30.9 | 1034.8 | ± | 114.5 |
| Fraction Photosynthetically Active Radiation (fPAR) | Fpar fraction | 49.2 | ± | 0.8 | 28.4 | ± | 0.6 | 26.0 | ± | 0.5 | 17.6 | ± | 0.6 |

| | | | | | |
|---|---|---|---|---|---|
| Soil water capacity at 5 cm depth | % | 22.6 ± 0.4 | 22.9 ± 0.3 | 27.0 ± 0.5 | 28.2 ± 0.2 |
| Northness | index (-1 to 1) | 0.1 ± 0.0 | 0.3 ± 0.0 | 0.0 ± 0.0 | -0.1 ± 0.0 |
| Potential Evapotranspiration (PET) | PET value (mm) | 987.5 ± 14.0 | 1305.1 ± 22.5 | 534.4 ± 5.7 | 655.6 ± 27.5 |
| Saturated Water Content 5 cm depth | % | 57.2 ± 0.5 | 53.0 ± 0.6 | 69.2 ± 0.4 | 63.1 ± 0.3 |
| Soil pH (water) at 5 cm depth | pH x 10 | 52.9 ± 0.5 | 68.3 ± 0.7 | 49.7 ± 0.3 | 61.0 ± 0.6 |

**Table S5** Results of single effects multivariate linear mixed-effects models for reported and measured macro-environmental factors with the standardised mean difference of decomposition (SMD) as dependent and measured or reported site-specific environmental factors as predictor. Values in bold indicate significant effect of the predictor on decomposition SMD (p ≤ 0.05). The number of effect sizes (*k*) used in the models, lower and upper bounds of the 95% confidence intervals, and heterogeneity explained by the model structure ($Q_M$) are reported.

| Predictor | *k* | slope | 95%CI | Test of Moderators (Qm, p-value) |
|---|---|---|---|---|
| Absolute Latitude | 637 | -0.003 | -0.01, 0.004 | 0.63, p = 0.429 |
| Duration of warming before experiment | 598 | -0.003 | -0.01, 0.02 | 0.12, p = 0.725 |
| Mesh size | 637 | -0.039 | -0.08, 0.003 | 3.29, p = 0.069 |
| Carbon to Nitrogen ratio | 428 | 0.001 | -0.00 ,0.00 | 0.25, p = 0.615 |
| Ambient decomposability (mass loss % $d^{-1}$) | 617 | -0.269 | -0.45, -0.09 | **8.97, p<0.001** |

**Table S6** Map-based macro-environmental results of single multivariate linear mixed-effects models with the standardised mean difference of decomposition (SMD) as dependent variable and the map-derived macro-environmental factors as predictor. Values in bold indicate significant effect of the predictor on decomposition SMD (p ≤ 0.05). The number of effect sizes (*k*) used in the models, lower and upper bounds of the 95% confidence intervals, and heterogeneity explained by the model structure ($Q_M$) are reported.

| Predictor | *k* | slope | 95%CI | Test of Moderators (Qm, p-value) |
|---|---|---|---|---|
| **Temperature** | | | | |
| Annual Mean Temperature | 635 | -0.005 | -0.02, 0.01 | 0.60, p = 0.439 |
| Max Temperature of Warmest Month | 635 | -0.004 | -0.02, 0.01 | 0.26, p = 0.611 |
| Air temperature isothermality | 635 | -0.005 | 0.02, 0.01 | 0.57, p = 0.449 |
| Mean Diurnal Range | 635 | 0.006 | 0.02, 0.04 | 0.10, p = 0.757 |
| Mean Temperature of Coldest Quarter | 635 | -0.007 | -0.02, 0.01 | 1.20, p = 0.274 |
| Mean Temperature of Driest Quarter | 635 | -0.006 | -0.01, 0.003 | 1.66, p = 0.198 |
| Mean Temperature of Warmest Quarter | 635 | -0.005 | -0.02, 0.01 | 0.43, p = 0.512 |
| Mean Temperature of Wettest Quarter | 635 | 0.000 | - 0.01, 0.01 | 0.00, p = 0.965 |
| Min Temperature of Coldest Month | 635 | -0.007 | -0.02, 0.004 | 1.50, p = 0.222 |
| Annual Temperature Range | 635 | 0.005 | 0.01, 0.02 | 0.66, p = 0.418 |
| Temperature Seasonality | 635 | 0.000 | -0.00, 0.00 | 0.50, p = 0.478 |
| Mean Temperature during Incubation Period | 635 | -0.010 | -0.02, -0.00 | **3.91, p = 0.048** |
| **Precipitation** | | | | |
| Annual Precipitation | 635 | 0.000 | -0.00, 0.00 | 1.49, p = 0.237 |
| Precipitation of Coldest Quarter | 635 | 0.000 | -0.00, 0.001 | 0.07, p = 0.785 |
| Precipitation of Driest Month | 635 | 0.000 | 0.005, 0.005 | 0.00, p = 0.989 |
| Precipitation of Driest Quarter | 635 | 0.000 | -0.001, 0.001 | 0.01, p = 0.916 |
| Precipitation of Warmest Quarter | 635 | 0.001 | -0.00, 0.00 | 3.23, p = 0.072 |

| | | | | |
|---|---|---|---|---|
| Precipitation of Wettest Month | 635 | 0.002 | -0.00, 0.003 | 3.82, p = 0.051 |
| Precipitation of Wettest Quarter | 635 | 0.001 | -0.00, 0.001 | 3.34, p = 0.068 |
| Precipitation Seasonality | 635 | 0.003 | -0.00, 0.01 | 2.81, p = 0.094 |
| Sum Precipitation during Incubation Period | 629 | 0.000 | -0.00, 0.00 | 3.76, p = 0.053 |
| **Soil** | | | | |
| Bulk density at 5 cm depth | 635 | 0.000 | -0.001, 0.00 | 0.47, p = 0.495 |
| SOC Content at 5 cm depth | 635 | 0.000 | -0.002, 0.001 | 0.18, p = 0.670 |
| SOC Density at 5 cm depth | 635 | 0.000 | -0.00, 0.001 | 0.96, p = 0.328 |
| SOC Stock 0-5 cm depth | 635 | 0.000 | -0.01, 0.01 | 0.00, p = 0.947 |
| Sum of Total Nitrogen at 5 cm depth | 608 | 0.000 | -0.00, 0.00 | 0.04, p = 0.837 |
| Sum of Total Nitrogen at 15 cm depth | 608 | 0.000 | -0.00, 0.00 | 0.20, p = 0.658 |
| Sum of Total Nitrogen at 30 cm depth | 608 | 0.000 | -0.00, 0.00 | 0.11, p = 0.739 |
| **Other** | | | | |
| Annal Mean Solar Radiation | 635 | 0.000 | -0.00, 0.00 | 0.35, p = 0.553 |
| Aridity Index | 635 | 0.000 | -0.00, 0.00 | 1.24, p = 0.265 |
| Aspect Cosine | 635 | -0.026 | -0.13, 0.08 | **0.21, p < 0.001** |
| Aspect Sine | 635 | -0.070 | -0.25, 0.11 | 0.58, p = 0.446 |
| Cover Barren | 635 | -0.001 | -0.01, 0.003 | 0.57, p = 0.451 |
| Cover Cultivated | 635 | -0.001 | -0.01, 0.004 | 0.04, p = 0.842 |
| Cover Deciduous Broadleaf Trees | 635 | 0.008 | 0.001, 0.01 | **5.65, p = 0.017** |
| Cover Evergreen Broadleaf Trees | 635 | -0.001 | -0.02, 0.02 | 0.02, p = 0.901 |
| Cover Evergreen Needleleaf Trees | 635 | 0.001 | -0.01, 0.01 | 0.07, p = 0.786 |
| Cover Herbaceous | 635 | 0.002 | -0.002, 0.01 | 0.51, p = 0.475 |
| Cover Regularly Flooded | 635 | -0.003 | -0.01, 0.01 | 0.41, p = 0.521 |
| Cover Shrubs | 635 | -0.008 | -0.01, -0.001 | **5.60, p = 0.019** |
| Eastness | 635 | -0.232 | -0.62, 0.16 | 1.38, p = 0.241 |
| Elevation | 635 | 0.000 | -0.00, 0.0001 | 2.63, p = 0.105 |
| Fraction Photosynthetically Active Radiation (fPAR) | 635 | 0.003 | 0.01, 0.01 | 0.44, p = 0.505 |
| Soil water capacity at 5 cm depth | 635 | -0.0027 | -0.02, 0.01 | 0.10, p = 0.751 |
| Northness | 635 | 0.076 | -0.14, 0.29 | 0.48, p = 0.489 |
| Potential Evapotranspiration (PET) | 635 | 0.000 | -0.00, 0.00 | 0.19, p = 0.667 |
| Saturated Water Content 5 cm depth | 635 | 0.005 | -0.01, 0.02 | 0.67, p = 0.414 |
| Soil pH (water) at 5 cm depth | 635 | 0.004 | -0.01, 0.015 | 0.52, p = 0.471 |

### Supporting Information

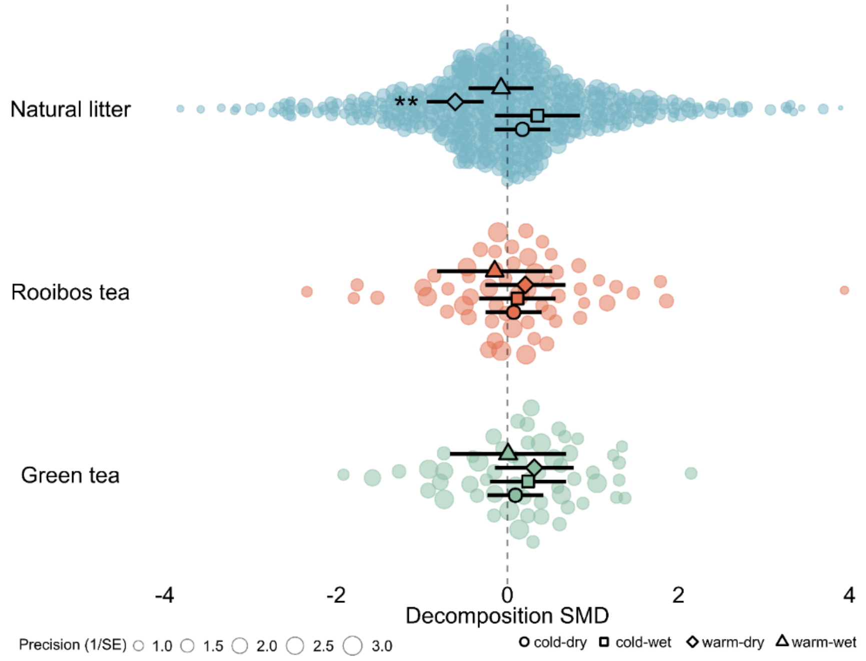


**Figure S3** Effects of experimental warming on plant litter decomposition. The pooled average decomposition standardised mean difference (SMD, Hedges' g; outlined circles) and 95% confidence intervals (black error bars) resulting from warming for the macro-environmental classes cold and dry (outlined circles), cold and wet (outlined squares), warm and dry (outlined diamonds), and warm and wet (outlined triangles) for the natural litter (blue, number of effect sizes k=523) and the standardised plant litter, separated into rooibos (red, k=57) and green tea (green, k=57). Each coloured dot is an individual effect size (non-outlined circles) with dot size representing its precision (the inverse of the standard error, larger points having greater influence on the model). Asterisks indicate that the overall pooled average SMD is significantly different from zero (**p < 0.01).

**Table S7** The impact of the four macro-environmental classes four macro-environmental classes distinguished by different combinations of high (▲) or low (▼) of temperature (temp), precipitation (prec) and soil organic carbon (SOC) and the natural and the standardised plant litter (i.e., green and rooibos tea) on the effect of warming on decomposition (SMD). Bold values indicate a significant effect of the macro-environmental class and litter type on SMD (p ≤ 0.05 or CI ≠ 0). Number of effect sizes (*k*), p-values, and 95%-confidence interval are shown.

| Macro-environment | litter type | SMD estimate | *k* | p-value | 95%CI |
|---|---|---|---|---|---|
| ▲ temp ▲ prec ▼ SOC | Natural litter | -0.07 | 155 | 0.703 | [-0.45; 0.30] |
| | Rooibos | -0.15 | 5 | 0.666 | [-0.82; 0.52] |
| | Green | 0.01 | 5 | 0.981 | [-0.67; 0.68] |
| ▲ temp ▼ prec ▼ SOC | Natural litter | **-0.61** | 150 | **<0.001** | [-0.94; -0.28] |
| | Rooibos | 0.21 | 10 | 0.382 | [-0.26; 0.68] |
| | Green | 0.31 | 10 | 0.180 | [-0.15; 0.77] |
| ▼ temp ▲ prec ▲ SOC | Natural litter | 0.35 | 126 | 0.167 | [-0.15; 0.85] |
| | Rooibos | 0.12 | 15 | 0.607 | [-0.33; 0.56] |
| | Green | 0.24 | 15 | 0.285 | [-0.20; 0.69] |
| ▼ temp ▼ prec ▲ SOC | Natural litter | 0.18 | 101 | 0.290 | [-0.15; 0.50] |
| | Rooibos | 0.07 | 27 | 0.659 | [-0.25; 0.40] |
| | Green | 0.09 | 15 | 0.575 | [-0.23; 0.42] |

#### Supporting Information

#### The effect of experimental-induced warming on decomposition

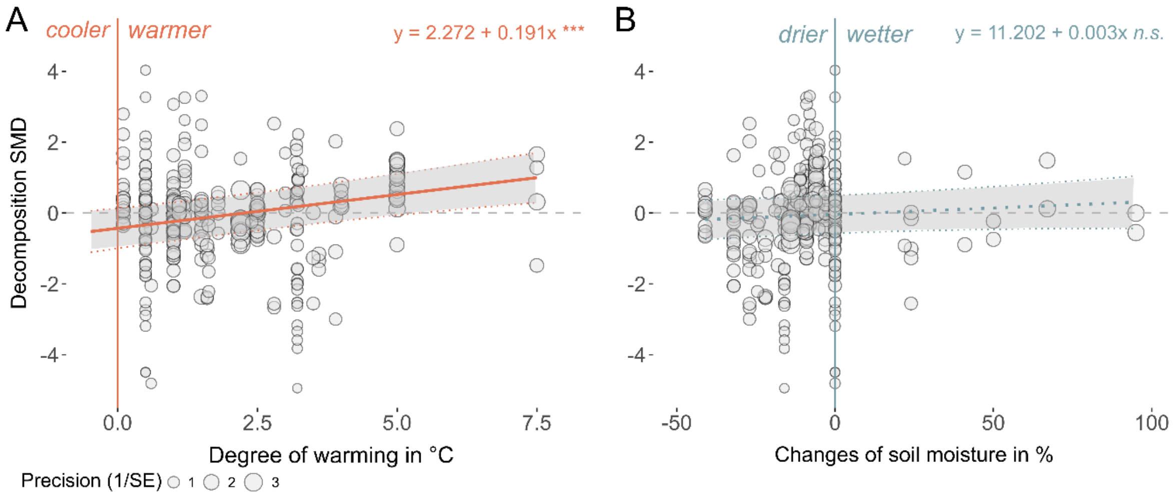


**Figure S4** Impacts of experimentally induced changes in micro-environment on decomposition and its interaction with macro-environment. Effect of **(A)** degree of warming (i.e., absolute temperature difference between warmed and control plots, k=315); and **(B)** warming-induced changes in soil moisture with warming (i.e., difference between warmed and control plots in soil moisture, k=315) on decomposition SMD. Each coloured dot is an individual effect size (non-outlined circles) with dot size representing its precision (the inverse of the standard error, larger points having greater influence on the model). Asterisks indicate that the overall pooled average SMD is significantly different from zero. Solid lines indicate regression lines with shaded areas representing the 95%CI (***p < 0.001). Dashed lines indicate no significant relationship (n.s. = not significant).

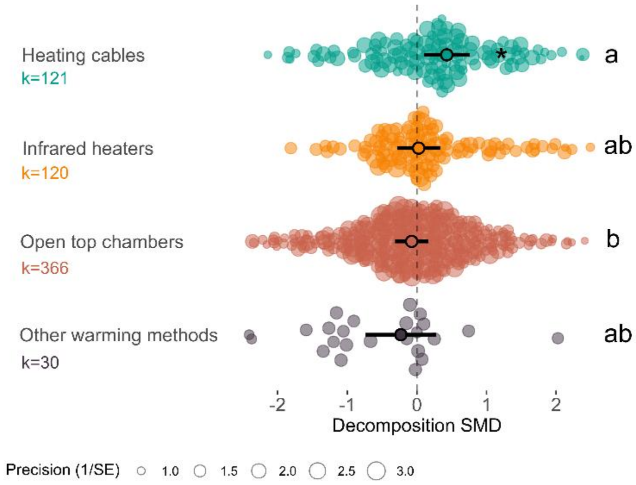


**Figure S5** Impact of warming methods on decomposition SMD. The pooled average decomposition standardised mean difference (SMD, Hedges' g; outlined circles) and 95% confidence intervals (black error bars) resulting from warming for the different experimental warming methods (see Table S1). Each coloured dot is an individual effect size (non-outlined circles) with dot size representing its precision (the inverse of the standard error, larger points having greater influence on the model). Letters indicate significant differences between the pooled average SMD of warming methods. Asterisks indicate a significant deviation of decomposition SMD from zero (*p ≤ 0.05).

### Supporting Information

Plant functional types and plant organ types interacting with the position of incubation (on soil surface, buried in the soil)

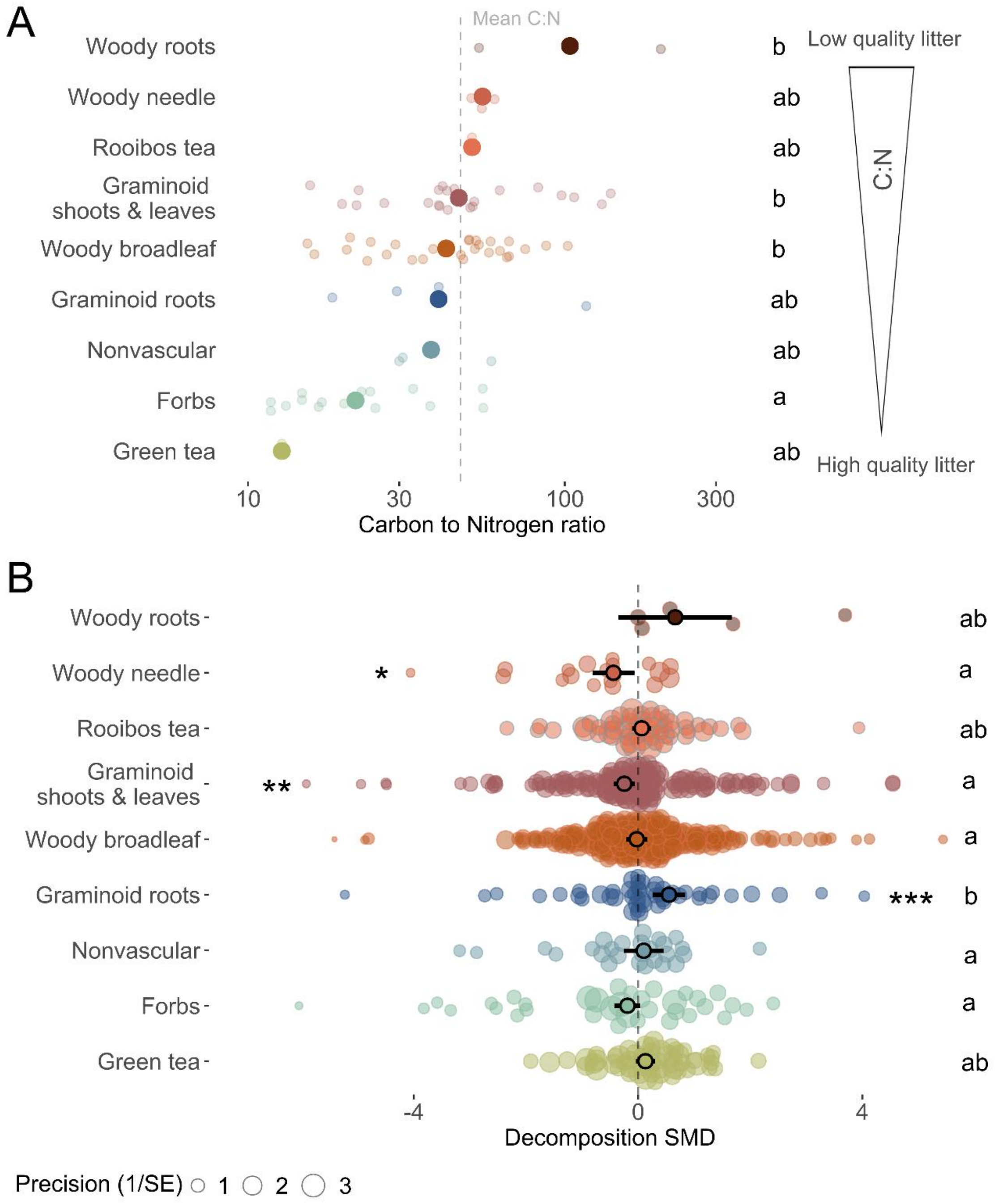


**Figure S6** Differences in C:N ratio and warming effect on decomposition across plant functional types**.** **(A)** Plant functional types ranked based on carbon to nitrogen ratios (C:N ratios). Large, coloured points represent mean C:N ratios and small transparent dots individual plant species. **(B)** The pooled average decomposition standardized mean difference (SMD, Hedges' g, black outlined circles) and 95% confidence intervals (95%CI, black error bars) per plant functional type of natural litter and standardised plant litter combining data from above and below ground incubations. Different letters indicate differences in **(A)** mean C:N ratio and **(B)** decomposition SMD between the different plant functional litter types, as well as the standard material green and rooibos tea. Asterisks indicate that the overall pooled average SMD is significantly different from zero (*$p < 0.05$, **$p < 0.01$, ***$p < 0.001$).

### Supporting Information

**Table S8** The pooled average decomposition standardised mean difference (SMD) of different plant functional types of the natural litter and natural and the standardised plant litter (i.e., green and rooibos tea) with respect to the position of incubation (i.e., on soil surface, buried in the soil) as well as the number of effect sizes (*k*) for each category, the p-value and 95%-confidence interval describing whether the pooled average SMD significantly differs from zero (in bold, p ≤ 0.05). For forbs and nonvascular plants no reports of buried or root litter were available.

| Plant functional type | Position incubated | *k* | SMD estimate | p-value | 95%CI |
|---|---|---|---|---|---|
| Forb | surface | 36 | -0.19 | 0.114 | [-0.42; 0.05] |
| Graminoid root | buried | 49 | **0.55** | **<0.001** | [0.27; 0.84] |
| Graminoid shoot/leaf | surface | 151 | **-0.25** | **0.010** | [-0.43; -0.06] |
| Green tea | buried | 57 | 0.13 | 0.133 | [-0.04; 0.30] |
| Nonvascular | surface | 27 | 0.10 | 0.589 | [-0.26; 0.45] |
| Rooibos tea | buried | 57 | 0.06 | 0.469 | [-0.11; 0.23] |
| Woody broadleaf | buried | 48 | -0.05 | 0.799 | [-0.44; 0.34] |
| Woody broadleaf | surface | 192 | -0.02 | 0.874 | [-0.21; 0.18] |
| Woody needle | surface | 21 | **-0.44** | **0.021** | [-0.82; -0.07] |
| Woody root | buried | 5 | 0.35 | 0.337 | [-0.37; 1.08] |

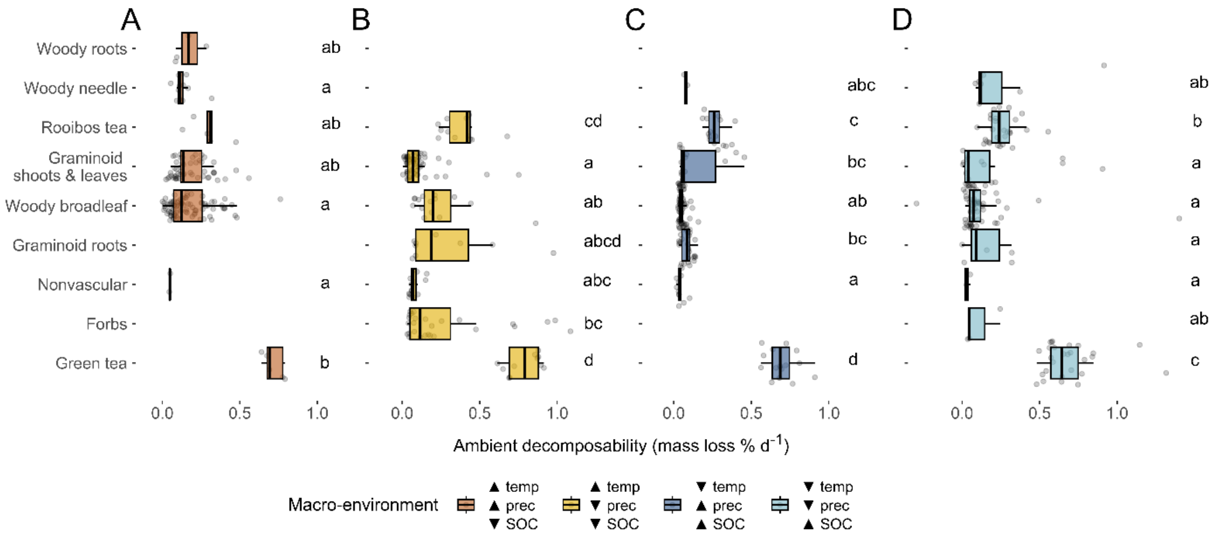


**Figure S7** Differences in ambient decomposability, measured as ambient mass loss rate per day (% $d^{-1}$), for the plant functional types and plant organs of natural plant litter and the standardised tea material (i.e., rooibos and green tea) for each of the four macro-environmental classes. Colours indicate the four macro-environmental classes of temperature (temp), precipitation (prec) and soil organic carbon (SOC) that are either high (▲) or low (▼), consistent with Figure 3 in the main text. Different letters indicate significant differences in decomposition SMD between plant functional types.
